# Mapping Neurophysiological Subtypes of Major Depressive Disorder Using Normative Models of the Functional Connectome

**DOI:** 10.1101/2023.02.13.528399

**Authors:** Xiaoyi Sun, Jin Liu, Qing Ma, Xiaoqin Wang, Dongtao Wei, Yuan Chen, Bangshan Liu, Chu-Chung Huang, Yanting Zheng, Yankun Wu, Taolin Chen, Yuqi Cheng, Xiufeng Xu, Qiyong Gong, Tianmei Si, Shijun Qiu, Ching-Po Lin, Jingliang Cheng, Yanqing Tang, Fei Wang, Jiang Qiu, Peng Xie, Lingjiang Li, Wenxu Wang, Yong He, DIDA-MDD Working Group, Mingrui Xia

## Abstract

Major depressive disorder (MDD) is the most burdensome psychiatric disorder characterized by remarkably heterogeneous clinical phenotypes. It remains challenging to delineate the heterogeneity of neurobiological abnormalities underlying the clinical variance and, on this basis, to identify neurophysiological subtypes of MDD patients. Here, using a large multisite resting-state functional MRI data from 1,148 MDD patients and 1,079 healthy controls, we generated lifespan normative models of functional connectivity strengths, mapped the heterogeneity of patients’ individual deviations, and identified neurobiological MDD subtypes. MDD patients showed positive deviations mainly in the default mode and subcortical areas, and negative deviations widely distributed over the cortex. However, there was a great inter-subject heterogeneity as indicated by that no more than 3.14% of patients deviated from the normative range for any brain region. Two neurophysiological MDD subtypes were identified. Subtype 1 showed severe deviations with positive deviations in the default mode, limbic, and subcortical areas, and negative deviations in the sensorimotor, dorsal and ventral attention areas, while subtype 2 showed a moderate but conversed deviation pattern. The severe-deviation subtype had older age, higher medicated proportion, and higher Suicide item score, while the moderate-deviation subtype showed higher Work and Activities and Depressed Mood item scores. Moreover, the baseline deviations in the severe-deviation subtype were predictive of 6-month antidepressant treatment effects in a subsample. To our knowledge, the current study is the largest multisite analysis of neurophysiological MDD subtyping to date and the findings shed light on our understanding of the biological mechanisms underlying the intersubject heterogeneity of clinical phenotypes, which are informative for the development of personalized treatments for this disorder.

## Introduction

Major depressive disorder (MDD) is one of the most prevalent and burdensome psychiatric disorders worldwide, and it is accompanied by heterogeneous emotional, neurovegetative, and neurocognitive symptoms [1, 2]. This clinical diversity among patients brings up a huge challenge for disease diagnosis and the prediction of course trajectories and treatment responses. However, the underlying neurophysiological substrates of this clinical heterogeneity remain largely unclear. Parsing the neurophysiological heterogeneity is essential to better link complex biological dysregulations with clinical manifestations, thus facilitating optimized treatment allocation for patients. Prior studies have attempted to identify MDD subtypes based on clinical symptoms, such as melancholic depression, atypical depression, and seasonal affective disorder [3-5]. These studies showed neurophysiological differences between the clinical subtypes and indicated a possible relationship between specific depressive symptom profiles and biological dysregulations. However, clinical symptoms interact in a complex manner with biological substrates and may change over age and disease course, the neurophysiological informed subtyping of MDD is still lacking. Exploring neurophysiological subtypes of MDD is expected to provide a more objective understanding of the biological mechanisms underlying the disorder and inform the development of personalized biomarkers for clinical diagnosis and treatment. This will help advance our understanding of the complex clinical heterogeneity of MDD and improve its diagnosis and treatment in the future.

Based on resting-state functional magnetic resonance imaging (r-fMRI), many case-control studies have documented the disrupted topological organization of the functional brain connectomes and identified several critical functional foci in MDD patients [6-9], which largely enhanced our understanding of the neurophysiological substrates of this disease. It is important to note that the results from the between-group comparisons in small-sample studies were largely inconsistent, and the effect size was small in recent large-sample multisite studies, which suggests a large degree of heterogeneity in functional connectome alterations in MDD patients. This has recently led to increase focus on the heterogeneity of functional connectomes in MDD patients [10-12], with growing attention on the investigation of neurophysiological subtypes based on functional connectomes [13-17]. Studies have found important roles for functional connectomes of default mode networks (DMN), limbic systems (LIM), and subcortical regions (SUB) in neurophysiological subtyping. For example, Liang *et al*. [15] found hyperconnectivity of DMN areas in one subtype and hypoconnectivity in the other subtype. Drysdale *et al*. [14] defined four neurophysiological subtypes based on the distinct functional connectivity patterns in LIM and frontostriatal networks. These studies observed differences in clinical presentations and treatment response among neurophysiological subtypes, which indicates the promise of discovering clinically valuable neurobiological subtypes based on functional connectomes. However, previous studies have largely ignored the fact that the functional connectomes can change dramatically over the lifespan and that individual abnormal measurements, obtained from a typical change, can provide more accurate and disease-specific information for subtyping. This aspect holds promise for the future personalized diagnosis and treatment for a more general population of MDD.

The normative model, a cutting-edge statistical framework that maps demographic or behavioral variables to a quantitative neuroimaging feature, has demonstrated its superiority in characterizing the expected change trajectory of neuroimaging features and identifying individual heterogeneous deviations from the norm [18-20]. Similar to the widely-used normative growth charts in pediatric medicine, where a child’s height or weight is compared to the normative distribution for that particular age and gender [21], the normative model can be used to evaluate individuals in relation to a neuroimaging normative feature at a particular age and gender. Recently, the normative model has gained increased attention in the field of psychiatric disorders, as it has been applied to characterize individual abnormalities and intersubject differences in neuroimaging features in disorders, such as autism [22-24], attention deficit/hyperactivity disorder [25], and schizophrenia [26]. Unlike the traditional case-control analysis that only provide information on group-level abnormities, the normative model takes into account intersubject differences within the patient and control groups and allows for measuring individual deviation from a large reference cohort. These individual deviations from the normative model are expected to complement the characterization of patients’ developmental abnormalities and aid in the detection of neurobiological subtypes with distinct biological dysregulations and clinical manifestations.

In this study, we conducted a comprehensive investigation into the neurobiological heterogeneity and subtypes of MDD using a large multisite r-fMRI dataset of 1,148 patients with MDD and 1,079 matched healthy controls (HCs). We adopted a novel normative model framework, which allows us to estimate individual deviations from the lifespan trajectory of functional connectivity strength (FCS). Through the analysis of these deviations, we aimed to uncover the intersubject heterogeneity among patients with MDD and identify neurobiological subtypes based on their deviation patterns. The identified neurobiological MDD subtypes were evaluated in the context of demographic and clinical variable differences.

## Materials and methods

### Imaging dataset and preprocessing

This study included 2,414 participants (1,276 patients with MDD and 1,138 HCs) from nine research centers through the Disease Imaging Data Archiving - Major Depressive Disorder Working Group (DIDA-MDD) [9]. All participants were diagnosed by experienced psychiatrists using structured clinical interviews. The patients met the Diagnostic and Statistical Manual of Mental Disorders-IV (DSM-IV) diagnostic criteria for MDD [27] and had no other Axis I disorder. The clinical symptoms of patients were assessed using the 17-item Hamilton Depression Rating Scale (HDRS-17). The HCs had no current or lifetime history of an Axis I disorder. After strict quality control for both clinical and imaging data (Supplement), the final sample consisted of 1,148 patients with MDD (aged 11-93) and 1,079 HCs (aged 13-81) (Table 1 and Supplementary Fig. 1). Additionally, a subsample of 43 patients (Supplementary Table 1) received a 6-month treatment with paroxetine (an antidepressant of selective serotonin reuptake inhibitor, SSRI), and the treatment outcomes were recorded (Supplement). This study was approved by the ethics committees of each research center. Written informed consent was obtained from all participants. All R-fMRI data of participants were obtained using 3.0-T MRI scanners. Detailed scanning parameters at each center are listed in Supplementary Table 2. R-fMRI data were then preprocessed using a standard pipeline as described in our previous work [9, 28] (Supplement).

**Table 1.**
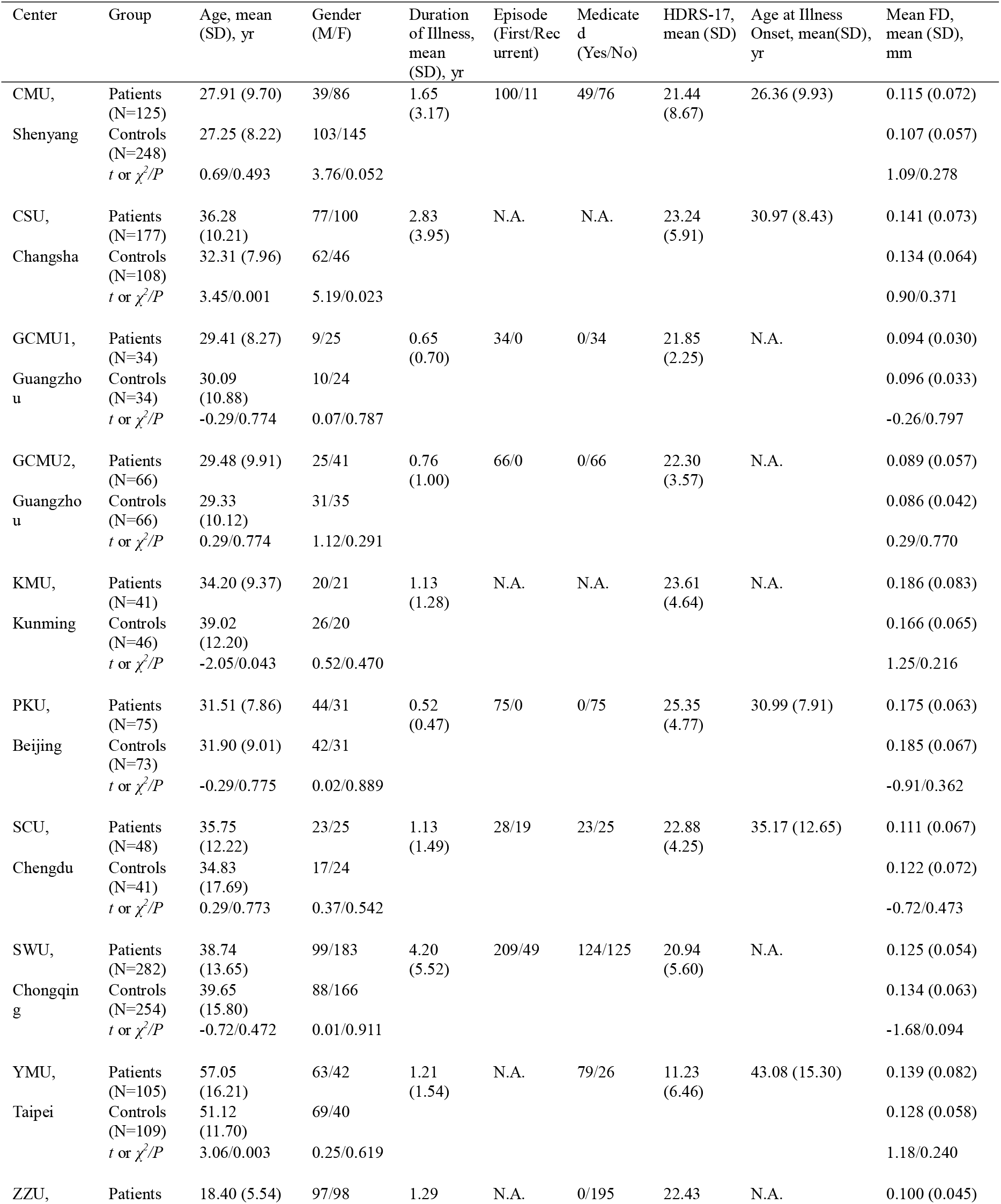
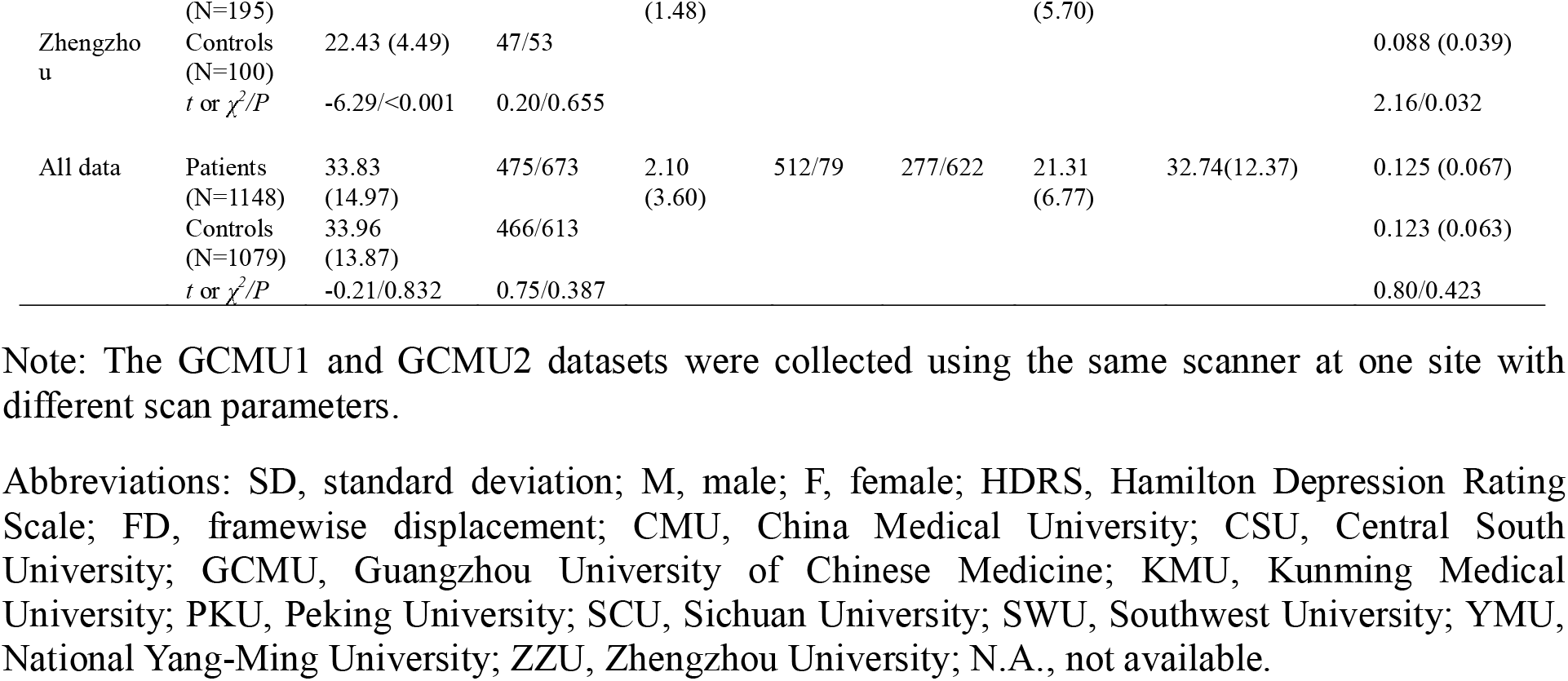
Demographic and clinical characteristics of the participants.

### Functional connectivity strength analysis

We first constructed a functional brain network for each participant. The network nodes were defined according to a predefined functional parcellation [29], including 220 cerebral regions that had qualified fMRI signals in all participants. The connectivity network was estimated by calculating Pearson’s correlation coefficients between the time series of any pairs of nodes followed by Fisher’s r-to-z transformation to improve normality. Then, the FCS values for each brain region was computed as the sum of the connectivity between a given region and all the other regions. Notably, we restricted our analysis to correlations above a threshold of r=0.2 to eliminate weak correlations possibly arising from noise, and the effects of different correlation thresholds on the results were validated (Supplement). The whole-brain FCS values were further standardized using z score normalization (minus the mean and divided by the standard deviation) to ensure comparability across participants. Finally, combat harmonization was utilized to correct the site effects on the FCS values [9, 28, 30-32].

### Normative modeling for functional connectivity strength

For each brain region, we estimated a normative model of FCS as a function of age and gender by using Gaussian process regression (GPR) [18] in the HCs (Fig. 1a and Supplement). GPR is a Bayesian nonparametric interpolation method that yields coherent measures of predictive confidence alongside point estimates [33]. In addition to fitting potentially nonlinear predictions of a brain feature, it can provide regional estimates of the expected variation in the relationship between age and brain features (normative variance) and estimates of uncertainty in this variance. The estimation of the normative models was performed using the PCNtoolkit package (https://github.com/amarquand/PCNtoolkit). To assess the generalizability of the models, we first estimated the normative models in the HCs under 10-fold cross-validation (Supplement), and overall standardized mean squared error and mean squared log-loss were used to evaluate the models. Then, the final normative models were trained on the whole HC dataset for the subsequent MDD deviation analyses.

**Fig. 1.**
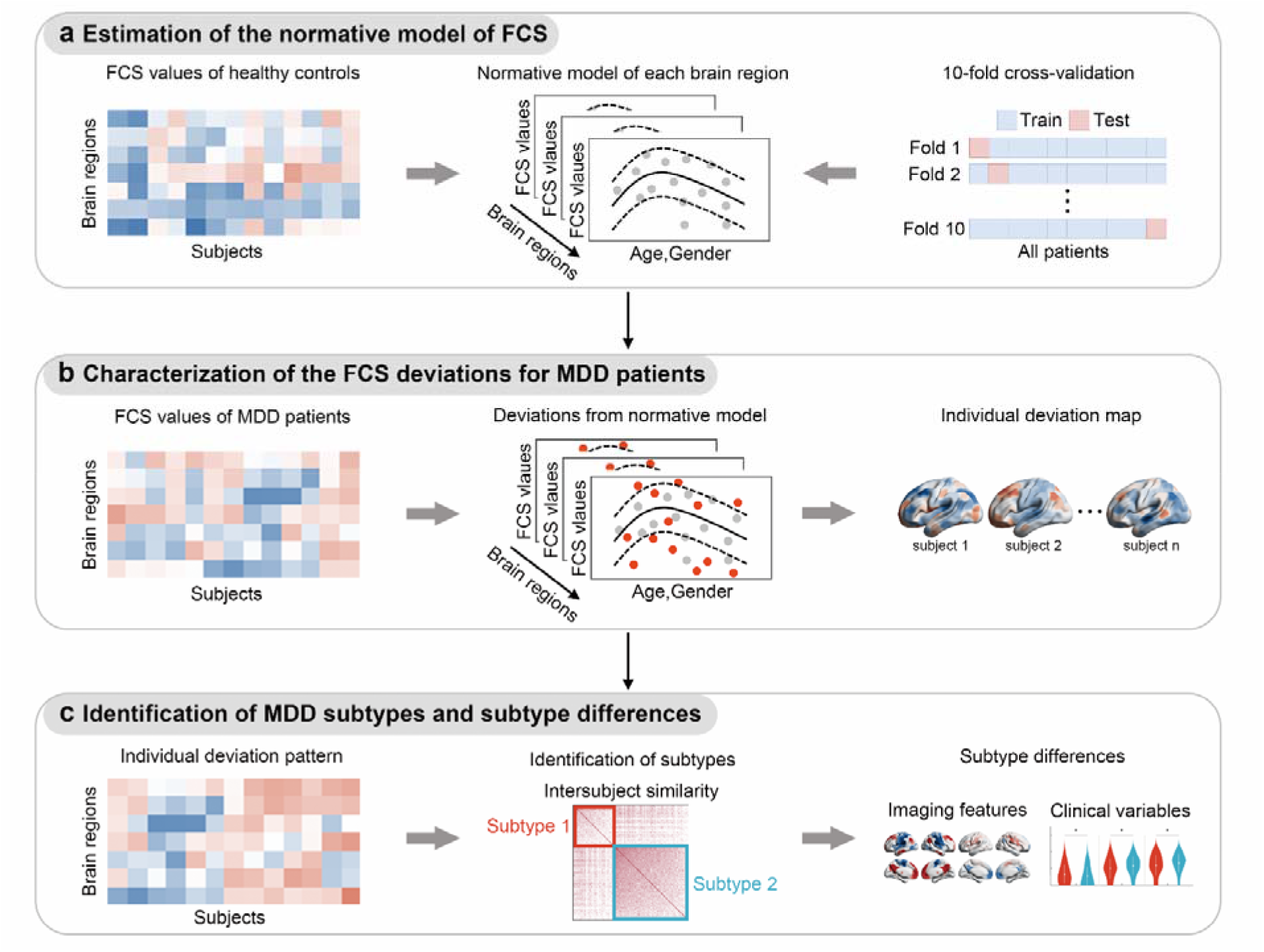
Flowchart of data analysis. **a** Estimation of the normative model of FCS for each brain region by training Gaussian process regression on the whole HCs dataset (gray dots). The solid line represents the predicted FCS values from the normative model, and the dashed line indicates the normative range. Ten-fold cross-validation were performed to assess the generalizability of the models. **b** Characterization of the FCS deviation of each brain region for each MDD patient (red dots) based on the normative model. **c** Identification of MDD subtypes based on the individual FCS deviation patterns and characterization of their imaging and clinical differences. GPR, Gaussian process regression; FCS, functional connectivity strengths; HCs, healthy controls; MDD, major depressive disorder.

### Estimating individual FCS deviations in normative models for MDD patients

For each patient with MDD, the FCS of the brain regions were positioned on the normative percentile charts from HCs to estimate individual deviation (Fig. 1b). We derived a *Z* value that quantifies the deviation from the normative model in each brain region [18]. For a given MDD patient *i*, the deviation *Z* value of a brain region *j* was calculated as follows:

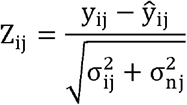

where y_ij_ is the observed FCS value, ŷ_ij_ is the predictive FCS value, σ_ij_ is the predictive uncertainty, and σ_nj_ is the variance learned from the normative distribution *n*. The *Z* value provides a statistical estimate of how much each patient differs from the healthy pattern in each brain region. Thus, the individual deviation map of each patient was obtained. The influence of patient sites on the calculation of FCS deviations was assessed in the validation (Supplement). Similarly, the individual deviation map of each HC participant was estimated by computing the *Z* value of each brain region during 10-fold cross-validation.

To further define the individual-level extreme deviations in the FCS of participants, we thresholded the deviation maps using *Z* = ± 2.6 (corresponding to a *p*<0.005) as was done in previous studies [25, 26, 34]. To quantify the overall extent of individual deviations, we calculated the number of brain regions with extreme deviations, the sum of positive extreme deviations, and the sum of negative extreme deviations for each participant. Then, to assess the intersubject heterogeneity of the deviations, we calculated a spatial overlap map by computing the percentage of participants who had an extreme deviation (*Z* > 2.6 or *Z* < -2.6) in each brain region. The between-group differences in the mean deviation map and the overall deviation indexes between patients with MDD and HCs were compared using two-sample t tests. The significance level was corrected for multiple comparisons using the FDR method (corrected *p*<0.05). The effect of different thresholds for defining extreme individual deviations on the results were validated (Supplement).

### Identifying MDD subtypes based on individual FCS deviations

We used a data-driven k-means clustering algorithm to explore MDD subtypes with different deviation patterns (Fig. 1c). The deviation map of each patient was set as the clustering feature, and the distance between any two patients was defined as the Euclidean distance between their deviation maps. The clustering algorithm was performed 10 times with different random initial cluster centroids to minimize the effect of the initial condition under each clustering number. The number of clusters was assessed from 2 to 10, and an optimal number of clusters was determined by a winner-take-all approach across 22 effective indexes using the NbClust package [35] (Supplement). To examine whether the MDD subtyping results were influenced by specific sites, we repeated the clustering analysis based on leave-one-site-out validation (Supplement).

### Characterizing subtype-related imaging and clinical differences

To investigate the deviation patterns between subtypes, we calculated the mean deviation map of each subgroup and compared them at the network level (Supplement). The overall deviated levels, including the number of extremely deviated regions, the sum of positive extreme deviations, and the sum of negative extreme deviations were compared among subtypes and HCs using one-way analysis of variance. Post hoc analysis was performed to compare the deviation differences between every two groups using two-sample t tests. To assess whether the deviated regions became more consistent after subtyping, we calculated the spatial overlap maps of extreme deviation for each subtype and compared them with the overlap maps of all patients using two-sample t tests (Supplement). The significance level was corrected for multiple comparisons using the FDR method (corrected *p*<0.05).

Group comparisons of demographic and clinical variables were performed on age, gender, disease duration, onset age, episode status, medication status, HDRS-17 total score, and HDRS-17 item scores using two-sample t tests or chi-square tests. Moreover, subtype differences in the association between the HDRS-17 total score and the duration/onset age were examined by using a one-way analysis of covariance (ANCOVA) with the duration/onset age as the predictor, the total HDRS-17 score as the response, and the subtypes as the grouping variable. The post hoc analysis was performed by calculating Pearson’s correlation coefficients between the HDRS-17 total score and the duration/onset age in each subtype. Support vector regression (SVR) was conducted to examine the prediction ability of deviation values for treatment response (i.e., changes in HDRS score) in patients. The baseline individual deviation values served as predictive features, and the model was validated using an embedded 5-fold cross-validation procedure and permutation tests (Supplement).

## Results

### Normative models of functional connectivity strength

The 10-fold cross-validation in the HCs revealed a high generalizability of the fitting performance of normative models for FCS, as indicated by overall standardized mean squared error close to 1 (0.996 ±0.013) and mean squared log-loss close to 0 (−0.001±0.007) (Supplementary Fig. 2). For the normative models established in the whole HCs, we found that the brain regions can be clustered (Supplement) into two categories according to their age-related FCS change trajectories in both female (Fig. 2a) and male groups (Supplementary Fig. 3). Specifically, regions with increased age-related FCS values were located mostly in the lateral frontoparietal cortices, dorsal anterior cingulate cortex, medial occipital cortices, sensorimotor areas, and subcortical areas, while those with decreased FCS were mainly in the precuneus, posterior cingulate cortex, medial prefrontal cortex, angular gyrus, insula, and medial temporal areas (Fig. 2a and Supplementary Fig. 3).

**Fig. 2.**
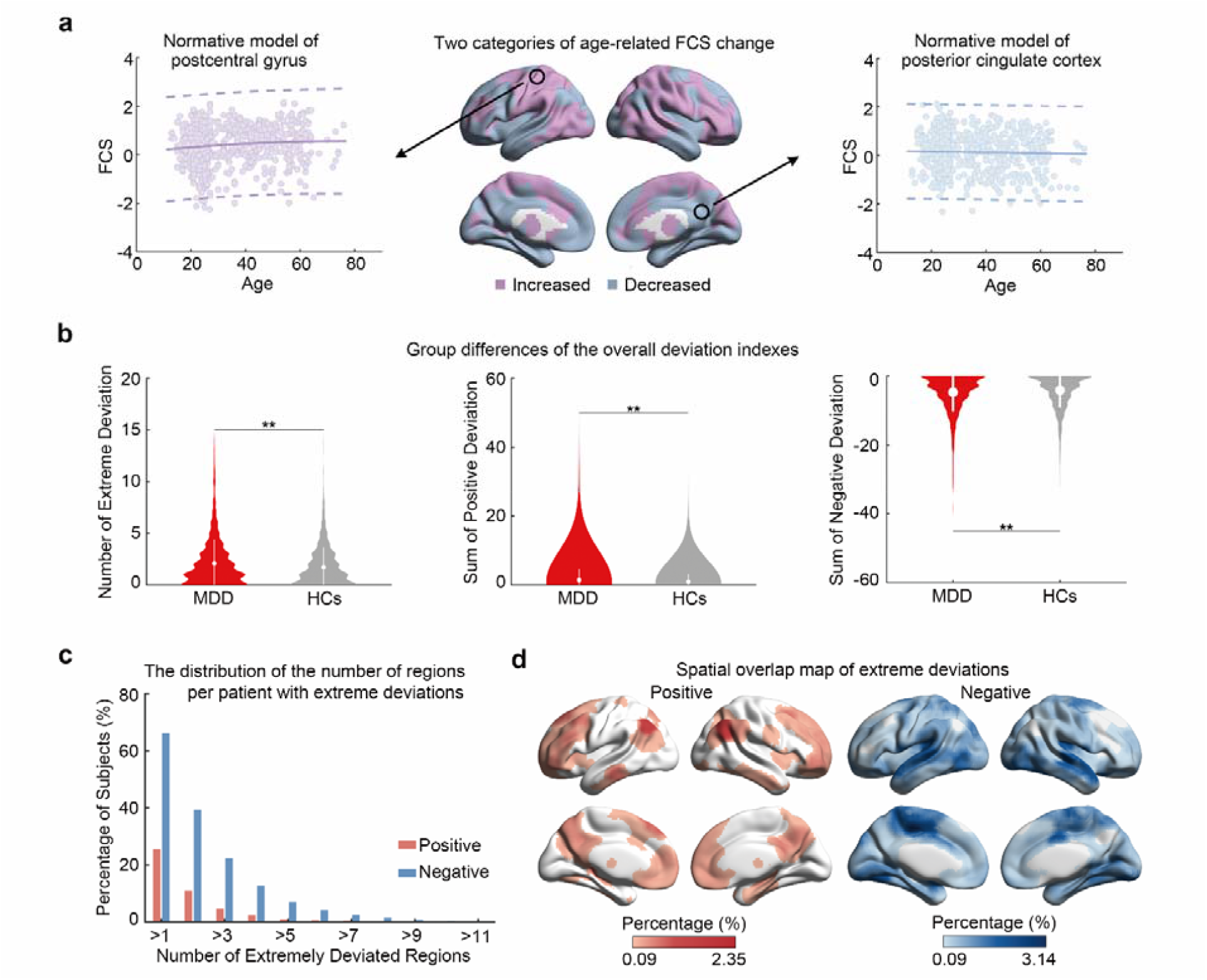
Normative models established in HCs and individual deviations from normative models in MDD patients. **a** The brain map in the middle indicates the two categories of age-related FCS change trajectories (purple: increased; blue: decreased) in HCs (female). The FCS change trajectories (solid line) and the normative range (dashed line) of postcentral gyrus and posterior cingulate cortex are shown on the left and right as examples. Each dot represents the data from one HC. **b** The between-group differences of the overall deviation indexes between patients with MDD and HCs. ^**^*p*<0.05, FDR corrected. **c** Bar plots show the distribution of the number of regions per patient with extremely positive (red) and negative (blue) deviations. **d** The spatial overlap maps indicate the percentage of patients who deviated extremely from the normative range for each brain region (left, extreme positive deviations; right, extreme negative deviations). FCS, functional connectivity strength; HCs, healthy controls; MDD, major depressive disorder.

### Highly heterogenous individual deviations from normative models in patients with MDD

Compared to the HCs, patients with MDD exhibited significantly larger individual FCS deviation indexes, including the number of extremely deviated regions (*t*=4.22) and the sum of positive (*t*=4.11) and negative (*t*=-2.77) extreme deviations (Fig. 2b, *p*<0.05, FDR corrected). Regionally, the patient group had significantly larger FCS deviations than the HC group, with positive deviations mainly in the bilateral lateral frontal cortex, precuneus, angular gyrus, and subcortical areas and negative deviations in the left parahippocampal gyrus, right Rolandic operculum, and middle cingulum gyrus (Supplementary Fig. 4 and Table 3, *p*<0.05, FDR corrected). A total of 72.82% (N=836) of the patients with MDD showed extreme FCS deviations from the normative model in at least one brain region, including extreme positive deviations in 25.78% (N=296) of patients and extreme negative deviations in 66.38% (N=762) of patients (Fig. 2c). From the perspective of brain regions, 99.55% (N=219) of the nodes showed an extreme FCS deviation in at least one patient, including extreme positive deviations in 67.73% (N=149) of brain regions and extreme negative deviations in 96.36% (N=212) of brain regions. The extreme positive deviations in patients with MDD were mostly located in the prefrontal cortex, precuneus, angular gyrus, and subcortical areas (Fig. 2d. left), and the extreme negative deviations were widespread over the whole brain, especially in the medial sensorimotor cortex and the temporal lobe (Fig. 2d. right). However, for any single brain region, the percentage of patients who deviated extremely from the normative range was remarkably low in either positive (≤2.35%, N=27) or negative (≤3.14%, N=36) deviations (Fig. 2d). These findings suggest that while alterations in FCS exist in most patients with MDD, the specific brain regions having out-of-range alterations varied remarkably among individual patients.

### FCS deviation-based MDD subtypes

The k-means clustering approach identified two MDD subtypes based on individual FCS deviations. This optimal subcluster number was consistently selected by 11 of 22 effective quality indexes (Fig. 3a). Patients with subtype 1 (37%, N=425) showed a severe deviation with positive deviations in the DMN, LIM, and SUB areas and negative deviations in the sensorimotor (SMN), dorsal attention (DAN), and ventral attention (VAN) areas (Fig. 3b and Supplementary Table 4, *p*<0.05, FDR corrected). However, the deviations observed in patients with subtype 2 (63%, N=723) were moderate, and the deviation patterns were significantly different, with negative deviations in the DMN, LIM, and SUB areas and positive deviations in the SMN, DAN, and VAN areas (Fig. 3b and Supplementary Table 4, *p*<0.05, FDR corrected). Statistical comparisons showed that the number of extremely deviated regions, the sum of positive extreme deviations, and the sum of negative extreme deviations observed in subtype 1 patients were significantly higher than those observed in HCs and subtype 2 patients, while the number of extremely deviated regions and the sum of negative extreme deviations observed in subtype 2 patients were significantly lower than those observed in HCs (Fig. 3c and Supplementary Table 5, *p*<0.05, FDR corrected). From the spatial overlap maps of extreme deviations, we observed a significantly higher consistency of extremely deviated regions among patients with the severe-deviation subtype compared to that among all patients (positive: 0.23-4.71%, *t*=3.31; negative: 0.23-5.88%, *t*=3.66; *p*<0.05, FDR corrected) and a significantly lower consistency among patients with the moderate-deviation subtype (positive: 0.13-2.49%, *t*=-3.26; negative: 0.13-1.80%, *t*=-3.79; *p*<0.05, FDR corrected) (Fig. 3d).

**Fig. 3.**
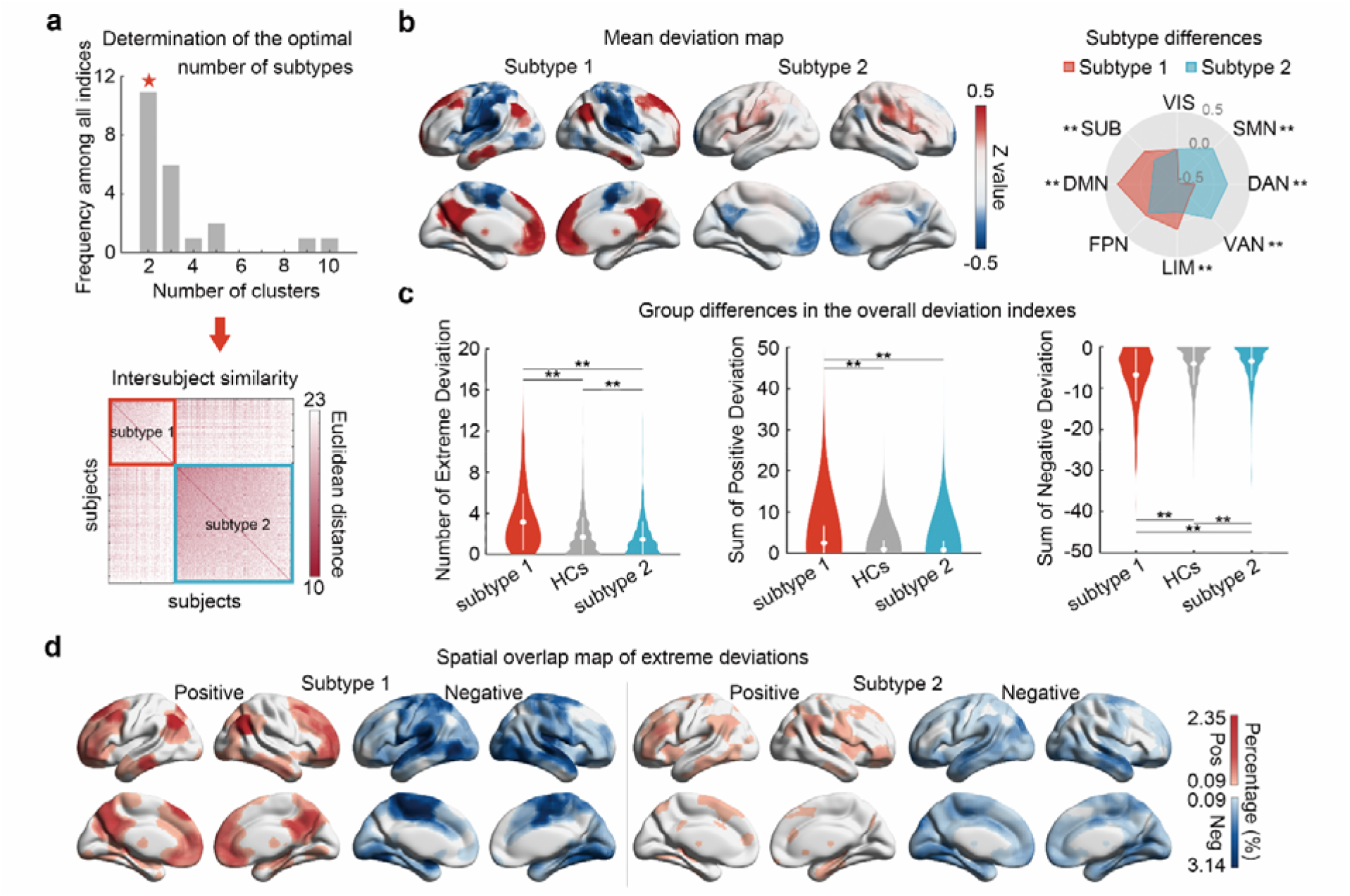
FCS deviation-based MDD subtypes. **a** Determination of the optimal number of MDD subtypes using the NbClust package and the intersubject similarity in the FCS deviation patterns among patients. **b** The mean deviation map of each subtype and their system-level differences. **c** The group differences in the overall deviation indexes among MDD subtypes and HCs. **d** The spatial overlap map of extreme positive and negative deviations of each subtype. VIS, visual network; SMN, sensorimotor network; DAN, dorsal attention network; VAN, ventral attention network; LIB, limbic network; FPN, frontoparietal network; DMN, default mode network; SUB, subcortical regions; HCs, healthy controls; MDD, major depressive disorder; ^**^*p*<0.05, FDR corrected.

Regarding demographic and clinical variables, patients with the severe-deviation subtype were significantly older (*t*=2.64, *p*=0.008) and had a higher medicated proportion (χ^*2*^=6.11, *p*=0.013) than patients with the moderate-deviation subtype (Fig. 4a and Supplementary Table 6). Patients with the severe-deviation subtype had more severe symptoms in the Suicide item (*t*=2.02, *p*=0.044), while patients with the moderate-deviation subtype exhibited more severe symptoms in the Work and Activities (*t*=3.11, *p*=0.002) and Depressed Mood items (*t*=2.42, *p*=0.016) (Fig. 4a and Supplementary Table 6). Moreover, ANCOVA showed that the correlations between the HDRS-17 score and the onset age were significantly different between the two subtypes (*F*=4.41, *p*=0.037) (Supplementary Table 7-8). The HDRS-17 score was negatively correlated with onset age in patients with the severe-deviation subtype (*r*=-0.24, *p*=0.004) but not in patients with the moderate-deviation subtype (*r*=-0.00, *p*=0.966) (Fig. 4b).

**Fig. 4.**
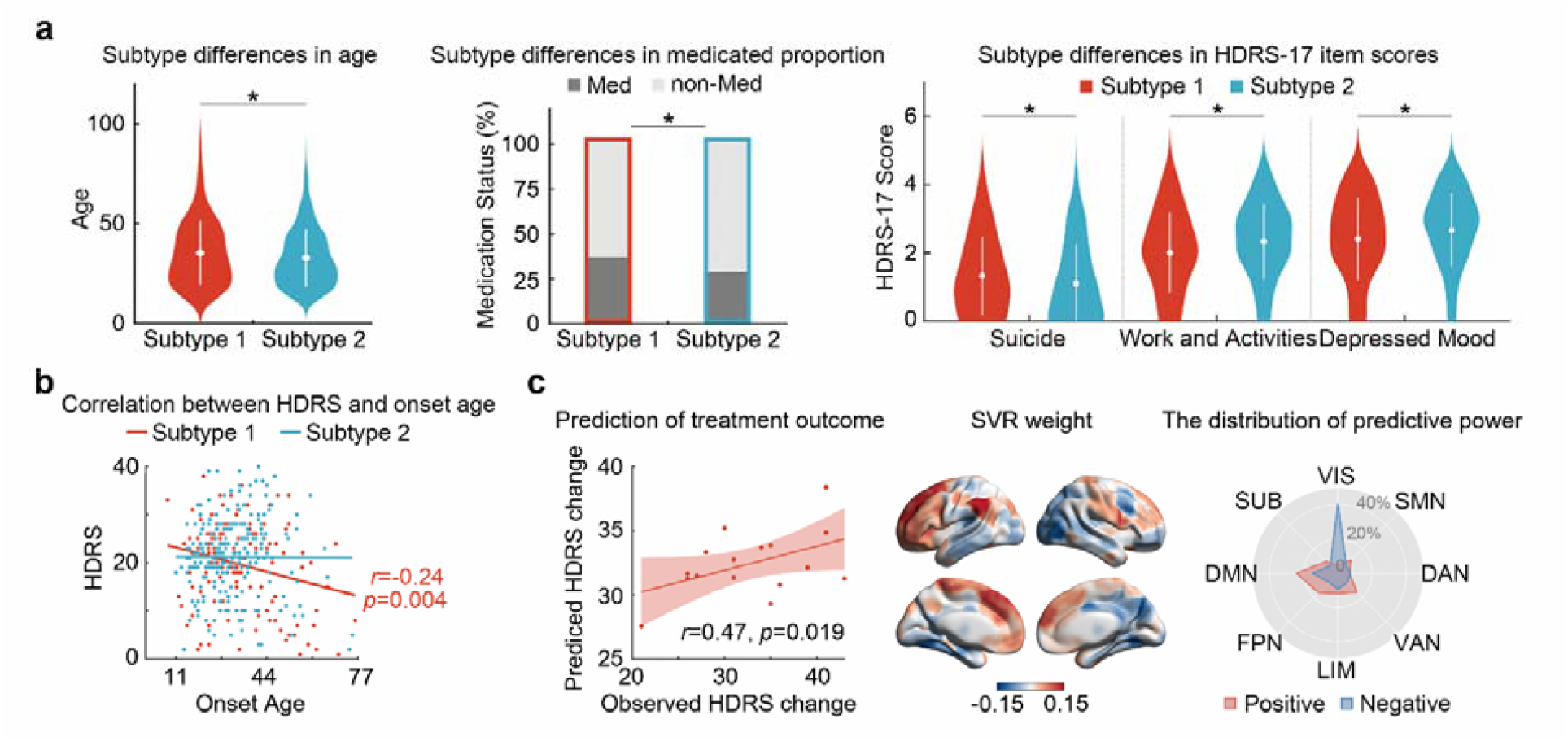
Subtype differences in demographic and clinical variables. **a** Subtype differences in age, medicated proportion, and HDRS-17 item score. ^*^*p*<0.05. **b** The correlation between the HDRS-17 total score and the onset age in each subtype. Each dot represents the data from one patient. **c** The prediction ability of deviation values for treatment response in patients of the severe-deviation subtype. The scatter plot presents the correlation between the observed HDRS score change after treatment and the predicted HDRS score change derived from the SVR. Each dot represents the data from one patient, and the dashes indicate the 95% prediction error bounds. The summed weights in 5-fold cross-validation were mapped onto the brain surface. The radar map represents the distribution of predictive power in different systems (red: positive; blue: negative). HDRS, Hamilton Depression Rating Scale; SVR, support vector regression; VIS, visual network; SMN, sensorimotor network; DAN, dorsal attention network; VAN, ventral attention network; LIB, limbic network; FPN, frontoparietal network; DMN, default mode network; SUB, subcortical regions.

Among the patients who had follow-up treatment outcomes, 16 patients were clustered into the severe-deviation subtype and the other 27 patients were clustered into the moderate-deviation subtype. The baseline individual deviation map could significantly predict HDRS score changes after treatment for patients with the severe-deviation subtype (*r*=0.47, *p*=0.019, one-tailed permutation test, Fig. 4c). The most positively contributive features were located in the DMN (24.1%), FPN (16.1%), and VAN (15.6%), and the most negatively contributive features were in the VIS (40.5%) (Fig. 4c).

In contrast, the baseline deviation map of the moderate-deviation subtype could not predict their HDRS score changes (*r*=-0.14, *p*=0.785, one-tailed permutation test).

### Validation results

Overall, the findings reported above were generally reproducible across different analytical choices. Under different thresholds in FCS calculation (r=0.15, 0.25), the normative models and patient deviations were similar to our main results, the overlap rates of the resulting subtype indexes with the clustered indexes in the main results were >96%, and the subtype differences largely remained (Supplementary Fig. 5-6 and Supplement). When different threshold was used to define extreme deviations (FDR *p*<0.05), the spatial overlap maps were slightly sparser, but the specific brain regions affected by MDD still varied markedly among individual patients (Supplementary Fig. 7). There were no significant site-related effects in the deviation values of all the brain regions (FDR *p*: 0.183∼0.100). The overlap rates of the resulting clustered indexes in the leave-one-site-out validation with the clustered indexes in the main results were all >92%, and the subtype differences were largely unchanged (Supplementary Fig. 8 and Table 9).

## Discussion

In this study, we uncovered the neurophysiological heterogeneity and subtypes of patients with MDD through mapping deviations from the normative models of functional connectome, by leveraging the currently largest R-fMRI dataset in MDD. Our findings reveal a significant intersubject variability in the spatial distribution of functional connectome abnormalities among MDD patients. Furthermore, our results highlight not only differences in the spatial distribution of functional connectome abnormalities but also significant disparities in demographic and clinical characteristics between the two identified neurobiological subtypes of MDD. Together, our study offers a novel analytical framework for subtyping MDD and offers promising implications for future personalized diagnosis and treatment of this disorder.

### Normative models of functional connectivity strength

Recently, several studies have estimated the normative model of brain structural features based on GPR, including cortical thickness, surface area, gray matter volume, white matter volume, and subcortical volume, and described the linear or nonlinear change trajectories of structures with age [23-26]. Compared to the traditional general linear model, the novel framework normative model does not require assumptions about the change trajectories and provides a useful tool to characterize any nonlinear changes in features. Here, based on a large sample dataset, we estimated the normative model of FCS for each brain region and found increased FCS values against age mostly in the lateral frontoparietal cortices, dorsal anterior cingulate cortex, medial occipital cortices, sensorimotor areas, and subcortical areas and decreased FCS mainly in the precuneus, posterior cingulate cortex, medial prefrontal cortex, angular gyrus, insula, and medial temporal areas. Similar to our findings, several previous studies found linear-age-related FCS decreases in the medial prefrontal cortex, precuneus, insula, and calcarine and linear FCS increases in sensorimotor areas based on the general linear model [36-38]. The areas of FCS decrease are the prominent hubs of global and local functional connectivity, and the age-related decrease could underlie the performance decline in working memory and visual sustained attention, which are the most affected cognitive functions that occur with aging [39-41]. Conversely, the sensorimotor areas are the least affected by aging [36]. These age-related changes in our study support the developmental theory which postulates that the first regions to emerge phylogenetically and ontogenetically are the most resistant to age effects, and the last ones are the most vulnerable. Notably, in our study, although brain regions had overall increased or decreased change trajectories, the changes did not always follow a linear or quadratic change, which demonstrates the high value of the normative model in characterizing the natural FCS change trajectories more accurately.

### Highly heterogenous individual deviations from normative models in MDD patients

In contrast to the case-control analysis identifying group-averaged alterations for patients, the normative model allows individual measures of the extent of patients’ deviation from a large reference cohort. Importantly, the model can recognize all sources of variance and reduce overly optimistic inferences and thus obtain more accurate and patient-specific individual deviations for patients. Given its great advantage, the normative model has recently been used to characterize the individual abnormalities and intersubject differences in neuroimaging features in multiple psychiatric disorders, such as autism [22-24], attention deficit/hyperactivity disorder [25], and schizophrenia [26]. Here, based on the normative model, our study investigated the individual FCS deviations for each patient and explored the heterogeneity of FCS deviations among patients. We found positive FCS deviations mainly in the DMN and SUB areas and negative deviations mainly in the sensorimotor and lateral temporal cortices. The FCS alterations in these regions have been proven to be related to the regulation of widespread cognitive, emotional, and executive control functions in patients with MDD [6, 42-50]. More importantly, we found that the overlap rates among patients in these regions were very low. This huge heterogenous among patients provides an important reference for the explanation of inconsistent results in prior functional connectome studies in MDD. For example, the medial prefrontal cortex showing heterogenous FCS alterations in our study was found to have both increased and decreased FCS in previous case-control studies [47, 51-53]. Our results suggest that FCS alteration is an important neuropathological feature of MDD, while the alteration patterns among patients are largely different and there might be multiple forms of MDD. Also, these findings reflect the useful application of normative model of functional connectome for identifying individual abnormities and parsing heterogeneity of MDD.

### FCS deviation-based MDD subtypes

Based on the individual FCS deviation pattern from normative models, we clustered MDD patients into two subtypes with distinct deviated levels and patterns. The FCS deviations in the DMN, LIM, SUB, SMN, DAN, and VAN exhibited significant differences and opposite alterations between the two subtypes, and the DMN showed the most. Consistent with the distinct DMN alterations in our study, several previous studies focusing on the local functional connectivity of the DMN or based on a small-sample dataset identified the different MDD subtypes with different functional connectivity patterns in the DMN areas [15, 54]. A transdiagnostic study, based on the whole brain amplitude of low-frequency fluctuations (ALFF), also clustered MDD patients into two subtypes with distinct activity patterns similar to our results [55]. Combined with these findings, our results indicate that the functional connectome and activity of DMN areas are important biomarkers for the neurophysiological subtyping of MDD. Among patients with MDD, there might be different disruption directions in DMN areas, some of them showed over-integration and increased activities in these areas, while the connectome and activities of these areas of other patients may not enough to support their normal functions. The different alteration patterns may result from complex genetic and environmental effects, which need to be further analyzed.

We found that the patients of the severe-deviation subtype showed more severe symptoms in the Suicide item score on the HDRS-17. Studies have shown that the increased functional connectomes and activities of the DMN and LIM areas are related to suicide, including the orbitofrontal cortex, medial prefrontal cortex, cingulate cortex, and striatum [56-59]. The orbitofrontal cortex is involved in learning, prediction, and decision-making for emotional and reward-related behaviors and is important in the regulation of behavioral impulsivity and response inhibition [60]. The higher FCS in the orbitofrontal cortex might be related to the increased vulnerability to suicidal behavior. The areas of the DMN are related to self-referential processing. Increasing evidence suggests that alterations in self-referential thinking may be associated with suicidal behavior [61]. When individuals are involved in regurgitating negative emotions about themselves, suicidal thoughts and behaviors occur in response to the individual’s desire to escape from both self-awareness and the associated unpleasant feelings [59]. On the other hand, the decreased functional connectomes in areas of the DMN and LIM are considered to be related to anhedonia [62-68], which is defined as diminished interest or pleasure in response to stimuli that were previously perceived as rewarding during a premorbid state [63]. Our results provide new evidence that the decreased FCS in the DMN and the LIM is related to the nonreactive mood and the failure to react to contextual changes in patients with MDD. More importantly, we found the predictive power of FCS deviation patterns for treatment effects in the severe-deviation subtype but not found in the other subtype. Studies have found that the recovery of increased DMN FCS has significant correlations with the treatment response [53], while decrease DMN FCS was associated with non-response to first-line antidepressants [15]. Together, the neurophysiological subtypes in our studies illuminated the different mechanisms underlying different clinical profiles and treatment responses among patients.

Patients with the severe-deviation subtype were older than patients with the moderate-deviation subtype. Previous studies have found different alteration patterns between patients in different age stages. Similar to the alteration patterns in our results, studies of late-life depression showed increased FCS in the inferior parietal lobule and parahippocampal gyrus and decreased FCS in the somatosensory and motor cortices compared to HCs [69]. The alterations in these areas might be related to the increased negative self-focused thought, impaired visuospatial and episodic memory, poor sleep quality, and deficits in physical health and functions in late-life patients. Moreover, evidence has shown that brain resilience increases during development and early adulthood and then decreases during aging [70]. Thus, the ability of the brain to withstand disease may decrease and thus experience more severe alterations in older patients. Additionally, a significant negative correlation between the onset age and HDRS-17 score was found only in the severe-deviation subtype. Several studies have explored the association between the onset age and HDRS-17 score in patient with MDD, but the results were inconsistent [71-74]. Our results indicated that these inconsistent observations may be contributed by different patient subtypes. Notably, in line with our findings, a study found that the onset age was negatively correlated with the cognitive-behavioral cluster of HDRS (including Suicide and Guilt item scores) but not with the affective cluster of HDRS (including Depressed Mood and Work and Activities item scores) [72]. In the severe-deviation subtype, the early-onset patients may disrupt the normal brain maturation, and thus leading to more severe symptoms of Suicide item. In the moderate-deviation subtype, the FCS alterations might be related to the higher symptoms in the Work and Activities and Depressed Mood items, which have lower effects with onset age.

### Limitations and future directions

Several issues with the current study need to be further addressed. First, our analysis was performed based on a cross-sectional sample. The age-related change trajectories shown here do not represent the trajectories of each participant, and they reveal age-specific population-level means and individual variabilities. Adding longitudinal samples will improve the representativeness of the brain change curve models. Second, in this study, we compared the subtype differences in clinical symptoms using HDRS-17 item scores. The patients with MDD also had varied cognitive impairments, which were not collected in the current retrospective study. Further analysis combined with more detailed cognitive performances could help us to better understand the complex relationship between the neurophysiological basis and the clinical presentations of MDD. Third, all the patients who were included in the analysis to predict treatment outcomes were responders to paroxetine, given that patients who had a poor response discontinued the medication or changed their treatment plans. Future studies need to include more nonresponders to establish prediction models for treatment-resistant depression and thus explore the different neuroimaging biomarkers between patients with different treatment outcomes. Finally, we identified MDD subtypes based on the heterogenous FCS alteration patterns of patients. An episode of MDD may be caused by numerous different factors, such as genetic liability, childhood adversity, and life stress [75-77]. Future studies combined with more genetic and environmental information are needed to investigate the factors that lead to the different neurophysiological subtypes.

## Acknowledgments

This work was supported by the SIT 2030-Major Projects (2022ZD0211500 to MX), National Natural Science Foundation of China (82071998 and 82021004 to MX; 81920108019, 91649117, 81771344, and 81471251 to SQ), Beijing Nova Program (Z191100001119023 to MX), Beijing United Imaging Research Institute of Intelligent Imaging Foundation (CRIBJZD202102 to MX), Science and Technology Plan Project of Guangzhou (2018-1002-SF-0442 to SQ), Guangzhou Key Laboratory (09002344 to SQ).

## DIDA-MDD Working Group

Yong He ^1,2,3,21^, Lingjiang Li ^9,10^, Jingliang Cheng ^8^, Qiyong Gong ^14,16^, Ching-Po Lin ^5,17^, Jiang Qiu ^6,7^, Shijun Qiu ^12^, Tianmei Si ^13^, Yanqing Tang ^18^, Fei Wang ^18^, Peng Xie ^19,20^, Xiufeng Xu ^15^ and Mingrui Xia ^1,2,3^

