## Supplement for "Mapping Neurophysiological Subtypes of Major Depressive Disorder Using Normative Models of the Functional Connectome"

SI Appendix for “Mapping Neurophysiological Subtypes of Major Depressive Disorder Using Normative Models of the Functional Connectome”: Supplemental Methods, Results, Tables, and Figures

SI Materials and methods

Imaging dataset and preprocessing

***Participants***

In this study, 1,276 patients with major depressive disorder (MDD) and 1,138 healthy controls (HCs) were recruited from nine research centers in China, including China Medical University (CMU), Central South University (CSU), Guangzhou University of Chinese Medicine (GCMU, two datasets), Kunming Medical University (KMU), Peking University Sixth Hospital (PKU), Sichuan University (SCU), Southwest University (SWU), National Yang-Ming University (YMU), and Zhengzhou University (ZZU). The patients met the Diagnostic and Statistical Manual of Mental Disorders-IV (DSM-IV) diagnostic criteria for MDD [1] and had no other Axis I disorder. The clinical symptoms of patients were assessed using the 17-item Hamilton Depression Rating Scale (HDRS-17). Disease duration, medication status, episode status, and onset age were collected for each patient. The HCs did not have a current or lifetime history of any Axis I disorder. The exclusion criteria for all participants included MRI contraindications, a history of drug or alcohol abuse, concomitant major medical disorder, a history of head trauma with consciousness disturbances, or any neurological disorders. Through strict quality control on clinical and imaging data, 187 participants were excluded due to a lack of demographic information (N=5), a young age (7 years old, N=1), a change in diagnosis during the follow-up interviews (N=7), duplicate data in the data transfer or errors in the raw DICOM data (N=17), different scanning parameters or incomplete scans (N=24), abnormalities in the anatomical brain images (N=14), excessive head motion (exceeding 3 mm of translational movement, 3° of rotational movement or 0.5 mm of mean framewise displacement, N=71), incomplete coverage of the entire brain (N=46), error in normalization (N=1), and an abnormal temporal signal-to-noise ratio (N=1). Ultimately, 2,227 participants (1,148 patients with MDD and 1,079 HCs) were included in the analysis. This study was approved by the ethics committees of each research center. Written informed consent was obtained from all participants.

In this dataset, 43 patients recruited from CSU (Supplementary Table 1) received a 6-month treatment with paroxetine (an antidepressant of the selective serotonin reuptake inhibitor, SSRI), and the treatment outcomes were recorded. Specifically, the patients were recruited from the inpatient or outpatient departments of the Zhumadian Psychiatry Hospital, Henan Province, China. Two trained psychiatrists confirmed the MDD diagnosis according to the DSM-IV criteria for MDD. Inclusion criteria for these MDD patients were as follows: acutely depressed and medication-free for not less than 2 weeks, and a score of at least 20 on the 24-item Hamilton Depression Rating Scale (HDRS-24). Exclusion criteria for patients included a comorbid Axis I or Axis II disorder or a personal history of bipolar disorder, substance abuse or dependence, neurological or internal illness, or any contraindication for MRI scans. All patients gave both written and verbal informed consent. This study was approved by the Human Investigation Committees of the Zhumadian Psychiatry Hospital and the Second Xiangya Hospital, Central South University of China. All patients were scanned at baseline and received a 6-month course of paroxetine treatment based on the judgment of the physician and the patient’s consent. In the first week, patients received 10 mg of paroxetine, which was the minimum dosing level for this study. In the second week, patients received 20 mg of paroxetine or higher dose levels depending on illness symptoms, clinical antidepressant effects, and side effects. The maximum dose was 60 mg of paroxetine. Symptoms of depression were assessed using the HDRS-24 in the sixth month.

***Data acquisition and preprocessing***

All R-fMRI data were obtained using 3.0-T MRI scanners. The participants were instructed to rest and relax with their eyes closed but to avoid falling asleep during scanning. The detailed scanning parameters at each center are listed in Supplementary Table 2. The R-fMRI data were preprocessed using SPM12 ([www.fil.ion.ucl.ac.uk/spm/](http://www.fil.ion.ucl.ac.uk/spm/)) and SeeCAT ([www.nitrc.org/projects/seecat](http://www.nitrc.org/projects/seecat)). Briefly, the first ten time points (the first five time points in the CSU, GCMU1, and ZZU datasets due to their short scan times) were discarded to allow scan equilibrium. The subsequent preprocessing steps included slice-timing correction and head-motion correction. The motion-corrected functional images were then normalized to the standard space using an echo planar imaging (EPI) template, resampled to 3-mm isotropic voxels, spatially smoothed with a 6-mm full-width at half-maximum Gaussian kernel, and linearly detrended. Several confounding covariates, including the Friston-24 head-motion parameters and the white matter and cerebrospinal fluid signals, were regressed from the time series for all voxels. Temporal bandpass filtering (0.01-0.1 Hz) was further applied. Finally, a “scrubbing” procedure was performed on the individual preprocessed datasets to remove outlier data due to head motion [2]. Specifically, volumes with a framewise displacement exceeding a threshold of 0.5 mm and their adjacent volumes (2 forward and 1 backward frames) were replaced with the linearly interpolated data.

Normative modeling for functional connectivity strength

For each brain region, we estimated a normative model of FCS as a function of age and gender by using Gaussian process regression (GPR). GPR is a Bayesian nonparametric interpolation method that yields coherent measures of predictive confidence alongside point estimates [3]. Briefly, a Gaussian process (GP) specifies a distribution over functions such that any finite number of elements has a joint Gaussian distribution. A GP can be denoted by $N\left( m\left( x \right),k(x,x') \right)$, which is specified by a mean $\left( m(x) \right)$ and covariance $\left( k(x,x') \right)$ function. Gaussian process models can be used for Bayesian nonlinear regression based on a set of training data $D=\left\{ x_{i},y_{i}\in\right\}_{i=1}^{N}$, where $x_{i}$ are the covariates (i.e.,: age, gender), $y_{i}$ are the response variables (i.e.,: FCS values at each brain region), and *N* is the number of participants. We use an unconstrained latent function ($f=\left[ f_{1},\ldots,f_{N} \right]^{T}$) to model the relationships between data points, which is assumed to differ from the true response variables by additive Gaussian noise, i.e.: $y_{i}=f_{i}+\varepsilon_{i}$ where $\varepsilon_{i}\sim N(0,\sigma_{n}^{2})$. The goal is to estimate this function from the training data in such a way that it allows us to accurately predict a new target $y_{*}$ from the test data $x_{*}$. To reach this end, a GP prior distribution is placed over the latent function and its posterior distribution is computed using the Bayes rule. A detailed description of the methods can also be found in Marquand *et al*. [4].

To assess the generalizability of the models, we first estimated the normative models in the HC group under 10-fold cross-validation. Specifically, we randomly partitioned the data into 10 folds. Nine folds of the data were used to train the model, and the remaining fold was used to estimate generalization performance. This procedure was repeated 10 times so that each fold was excluded once. We then trained the final normative models on the whole HC dataset for the following analyses.

After obtaining the normative models for predicting FCS trajectory against age and gender for each brain region, we applied a k-means clustering method to identify different categories of FCS change trajectories over the brain. The age-related FCS change trajectory of each brain region was set as the feature, and the distance between any two regions was defined as one minus the Pearson correlation coefficient between their FCS trajectories. The number of clusters was assessed from 2 to 10, and 10 iterations with different random initial cluster centroids were performed to minimize the effect of the initial condition under each clustering number. The optimal number of clusters was determined as the minimum value of the maximum mean silhouette value [5].

Identifying MDD subtypes based on individual FCS deviations

We used a data-driven k-means clustering algorithm to explore MDD subtypes with different deviation patterns. The deviation map of each patient was set as the clustering feature, and the distance between any two patients was defined as the Euclidean distance between their deviation maps. The number of clusters was assessed from 2 to 10 and an optimal number of clusters was determined by a winner-take-all approach across 22 effective indexes using the NbClust package [6]. All of the indexes were separately used to assess different aspects of the quality of the clustering, including the Krzanowski and Lai (KL) index, the Calinski and Harabasz (CH) index, the Hartigan index, the Cubic Clustering Criterion (CCC) index, the Marriot index, the Trcovw index, the Tracew index, the Friedman index, the Rubin index, the Cindex, the Davies and Bouldin (DB) index, the Silhouette index, the Duda index, the Pseudot2 index, the Beale index, the Ratkowsky index, the Ball index, the PtBiserial index, the Gap index, the Frey index, the McClain index, and the Gamma index.

Characterizing subtype-related imaging and clinical differences

To investigate the differences in deviation patterns between MDD subtypes, we computed the mean deviation map for each subtype and compared them on the network level. Specifically, we first obtained a template including eight networks: the visual, sensorimotor, dorsal attention, ventral attention, limbic, frontoparietal, and default mode networks defined by Yeo *et al.* [7] and the subcortical system including subcortical regions defined by the Automated Anatomical Labelling (AAL) atlas [8, 9]. The 220 brain regions in our analysis were assigned to the network with the most voxel attributions in that brain region. Then, we calculated the mean deviation value of each network for each patient. Two-sample t tests were used to compare the mean deviation values between subtypes at the network level. To assess whether the deviated regions became more consistent after subtyping, we calculated the spatial overlap maps of extreme positive and negative deviations for each subtype. The overlap values of each subtype were compared with those of all patients using two-sample t tests. The significance level was corrected for multiple comparisons using the FDR method (corrected *p*<0.05).

We performed support vector regression (SVR) with a linear kernel to examine the ability of the individual deviations to predict the improvement in symptoms (i.e., longitudinal changes in HDRS-24 total score) of antidepressant treatment in patients. A 5-fold cross-validation strategy was adopted to estimate the accuracy of the prediction. Here, the participants were first sorted according to their longitudinal changes in HDRS-24 score and were then assigned to the corresponding fold (e.g., 1^st^, 6^th^, 11^th^, …, to the first fold; 2^nd^, 7^th^, 12^th^, …, to the second fold; 3^rd^, 8^th^, 13^th^, …, to the third fold; 4^th^, 9^th^, 14^th^, …, to the fourth fold, and 5^th^, 10^th^, 15^th^, …, to the fifth fold). Four folds of the data were defined as the training set in turn, and the remaining fold was defined as the test set. During each training procedure, an internal 5-fold cross-validation was performed to determine the optimal parameters for relevant regression algorithms (i.e., C = 2^-5^, 2^-4^, …, 2^9^, 2^10^). To avoid the bias caused by features with greater numeric ranges, we z-normalized each feature in the training dataset and applied the estimated parameters to the testing dataset. The accuracy was reported as the Pearson’s correlation coefficient between the predicted and observed HDRS-24 score changes across all patients. The nonparametric permutation test (1,000 times) was performed to assess the statistical significance of the prediction accuracy. During each permutation, the patients’ observed HDRS-24 score changes were randomly shuffled before SVR and cross-validation. Thus, the null distribution of the correlation coefficients was obtained, and a *p* value was calculated by dividing the number of times that the permutations had higher correlation coefficients than the observed coefficient by 1,000. The codes for this prediction analysis were mainly conducted with libsvm (www.csie.ntu.edu.tw/~cjlin/libsvm/) and modified codes from *Cui and Gong* [10] (<https://github.com/ZaixuCui/Pattern_Regression_Matlab>). The system distribution of predicted weight was computed based on a 7-system cortical parcellation [7] and a subcortical system of the AAL atlas [8, 9]. Specifically, the system weights were calculated by summing the positive weight and negative weight of all brain regions belonging to this network. The proportion of each network was obtained by dividing the system weights by the total weight.

Validation analysis

To assess the reliability of our research, we calculated whole-brain FCS values on the different correlation thresholds, including r=0.15 and 0.25. The FCS values were further standardized using z score normalization, and the site effects were corrected using Combat harmonization. Based on these FCS values, we re-estimated the normative models for each brain region in the HC group, and the different categories of FCS change trajectories over the brain were identified based on the k-means clustering method. The resulting clustered indexes were compared with the clustered index in the main results by calculating the overlap rate. Then, the individual deviation patterns for patients with MDD were obtained based on the normative model. To assess the intersubject heterogeneity of the deviations, the number of extremely deviated regions of each patient and the spatial overlap map of extreme deviation among patients were explored. Finally, the MDD subtypes were reidentified based on the individual deviation patterns. The resulting clustered indexes were compared with the clustered index in the main results by calculating the overlap rate. The mean deviation maps and the spatial overlap maps of each subtype were calculated and compared. The demographic and clinical differences were recomputed between the resulting subtypes.

In the main text, we thresholded the deviation maps using Z=±2.6 and quantified the individual extreme deviations of patients with MDD. The fixed statistical threshold across participants allows for a simplified comparison between participants in terms of the number of extreme deviations from the normative model, even when the overall distribution of deviations of a participant is shifted [11]. We also performed the analyses again by thresholding the individual extreme deviations at an FDR-corrected *p*<0.05. This process controls for multiple comparisons across brain regions, while estimating a separate threshold for each participant. The spatial overlap maps of extreme deviation and the number of extreme deviations were re-examined.

To assess the influence of patient sites on the calculation of FCS deviations, for each brain region, we compared the deviation values among sites using one-way ANOVA. Multiple comparisons across brain regions were corrected (*p*<0.05, FDR corrected). To examine whether the MDD subtyping results were influenced by specific sites, we repeated the clustering analysis based on leave-one-site-out validation. Specifically, we performed a k-means analysis on deviation maps of patients excluding one site and calculated the overlap rate of the resulting clustered indexes with the clustered indexes in the main results. Demographic and clinical differences were also computed between the resulting subtypes. This procedure was repeated 10 times so that each site was excluded once.

SI Results

Validation results

For the normative models that were established based on the FCS that was calculated under the threshold of r=0.15, the age-related FCS change trajectories of brain regions were very similar to those in the main results. Specifically, the brain regions can be clustered into two categories according to their change trajectories, and the overlap rate between the new cluster indexes and the cluster indexes in our main results was higher than 95% (96.82% for female, 95.45% for male, Supplementary Fig. 5a). For the individual-level extreme deviations, 75.61% (N=868) of the patients with MDD patients showed extreme FCS deviations from the normative model in at least one brain region, including extreme positive deviations in 22.82% (N=262) of patients and extreme negative deviations in 71.17% (N=817) of patients (Supplementary Fig. 5b). From the perspective of brain regions, 99.55% (N=219) of the nodes showed an extreme FCS deviation in at least one patient, including extreme positive deviations in 65.00% (N=143) of brain regions and extreme negative deviations in 95.91% (N=211) of brain regions. The brain areas of extreme positive and negative deviations were similar to the main results (Supplementary Fig. 5b), and the percentage of patients who deviated extremely from the normative range was remarkably low in positive (≤2.09%, N=24) and negative (≤3.22%, N=37) deviations. The k-means clustering approach identified two MDD subtypes based on individual FCS deviations, including 416 (36.24%) patients with subtype 1 and 732 (63.76%) patients with subtype 2. This optimal subcluster number was consistently selected by 10 of 22 effective quality indexes (Supplementary Fig. 5c). The overlap rate of the resulting clustered indexes with the clustered indexes in the main results was 99.22%. Patients with subtype 1 showed severe deviations, including positive deviations in the DMN, LIM, and SUB areas and negative deviations in the SMN, DAN, and VAN areas (*p*<0.05, FDR corrected) (Supplementary Fig. 5d). The deviations among patients with subtype 2 were moderate, and the deviation patterns were significantly different, with negative deviations in the DMN, LIM, and SUB areas and positive deviations in the SMN, DAN, and VAN areas (*p*<0.05, FDR corrected) (Supplementary Fig. 5d). From the spatial overlap maps of extreme deviations, we observed a significantly increased consistency of extremely deviated regions among patients of the severe-deviation subtype compared to that among all patients (positive: 0.24-4.33%, *t*=3.14; negative: 0.24-6.25%, *t*=3.88; *p*<0.05, FDR corrected), which were decreased among patients of the moderate-deviation subtype (positive: 0.14-2.05%, *t*=-3.07; negative: 0.13-2.05%, *t*=-3.92; *p*<0.05, FDR corrected) (Supplementary Fig. 5e). Regarding demographic and clinical variables, patients with the severe-deviation subtype were significantly older (*t*=2.40, *p*=0.017) and had a higher medicated proportion (*χ^2^*=5.29, *p*=0.021) compared with patients with the moderate-deviation subtype (Supplementary Fig. 5f). Patients of the moderate-deviation subtype exhibited more severe symptoms in the Work and Activities (*t*=3.15, *p*=0.002) and Depressed Mood item scores (*t*=2.10, *p*=0.036) (Supplementary Fig. 5f). The Suicide item and the association between HDRS-17 score and onset age between the two subtypes had a trend towards differences (*t*=1.85, *p*=0.065; *F*=3.64, *p*=0.057). Among the patients who had follow-up treatment outcomes, 16 patients were clustered into the severe-deviation subtype and the other 27 patients belonged to the moderate-deviation subtype. The overlap rate of the resulting clustered indexes with the clustered indexes in the main results was 100%.

For the normative models that were established based on the FCS that was calculated under the threshold of r=0.25, the age-related FCS change trajectories of brain regions were also very similar to those in the main results. Specifically, the brain regions can be clustered into two categories according to their change trajectories, and the overlap rate between the new cluster indexes and the cluster indexes in our main results was 97.73% for both female and male participants (Supplementary Fig. 6a). For the individual-level extreme deviations, 70.21% (N=806) of the patients with MDD showed extreme FCS deviations from the normative model in at least one brain region, including extreme positive deviations in 31.36% (N=360) of patients and extreme negative deviations in 58.89% (N=676) of patients (Supplementary Fig. 6b). From the perspective of brain regions, 99.55% (N=219) of the nodes showed an extreme FCS deviation in at least one patient, including extreme positive deviations in 73.00% (N=162) of brain regions and extreme negative deviations in 92.73% (N=204) of brain regions. The brain areas of extreme positive and negative deviations were similar to those in the main results (Supplementary Fig. 6b), and the percentage of patients who deviated extremely from the normative range was remarkably low in positive (≤2.79%, N=32) and negative (≤3.05%, N=35) deviations. The k-means clustering approach identified two MDD subtypes based on individual FCS deviations, including 425 (37.02%) patients with subtype 1 and 723 (62.98%) patients with subtype 2. This optimal subcluster number was consistently selected by 9 of 22 effective quality indexes (Supplementary Fig. 6c). The overlap rate of the resulting clustered indexes with the clustered indexes in the main results was 96.34%. Patients with subtype 1 showed a severe deviation with positive deviations in the DMN, LIM, and SUB areas and negative deviations in the SMN, DAN, and VAN areas (*p*<0.05, FDR corrected) (Supplementary Fig. 6d). The deviations of patients of subtype 2 were moderate, and the deviation patterns were significantly different, with negative deviations in the DMN, LIM, and SUB areas and positive deviations in the SMN, DAN, and VAN areas (*p*<0.05, FDR corrected) (Supplementary Fig. 6d). From the spatial overlap maps of extreme deviations, we observed a significantly increased consistency of extremely deviated regions among patients of the severe-deviation subtype compared to that among all patients (positive: 0.24-4.71%, *t*=3.34; negative: 0.24-5.41%, *t*=3.48; *p*<0.05, FDR corrected), which were decreased among patients of the moderate-deviation subtype (positive: 0.14-2.77%, *t*=-3.21; negative: 0.14-1.66%, *t*=-3.65; *p*<0.05, FDR corrected) (Supplementary Fig. 6e). Regarding demographic and clinical variables, patients with the severe-deviation subtype had more severe symptoms in the Suicide item score (*t*=2.03, *p*=0.043), while patients with the moderate-deviation subtype exhibited more severe symptoms in the Work and Activities (*t*=2.80, *p*=0.005) and Depressed Mood item scores (*t*=2.26, *p*=0.024) (Supplementary Fig. 6f). Patients with the severe-deviation subtype were significantly older (*t*=2.64, *p*=0.008) and had a higher medicated proportion (*χ^2^*=6.11, *p*=0.013) than patients with the moderate-deviation subtype (Supplementary Fig. 6f). Moreover, ANCOVA showed that the correlations between HDRS-17 score and onset age were significantly different between the two subtypes (*F*=4.35, *p*=0.038). Post hoc analysis showed that the HDRS-17 score was negatively correlated with the onset age in patients with the severe-deviation subtype (*r*=-0.24, *p*=0.004), but not in patients with the moderate-deviation subtype (*r*=-0.00, *p*=0.966) (Supplementary Fig. 6f). Among the patients who had follow-up treatment outcomes, 17 patients were clustered into the severe-deviation subtype and the other 26 patients belonged to the moderate-deviation subtype. The overlap rate of the resulting clustered indexes with the clustered indexes in the main results was 97.67% (N=42), while the baseline deviation map of the severe-deviation subtype could not predict their HDRS changes as in the main results (*r*=0.15, *p*=0.274). Future studies are needed to include more patients to establish prediction models for treatment-resistant depression and thus explore neuroimaging biomarkers among patients with different treatment outcomes.

When we thresholded the deviation maps using an FDR-corrected *p*<0.05, 5.57% (N=64) of the patients with MDD showed extreme FCS deviations from the normative model in at least one brain region, including extreme positive deviations in 0.96% (N=11) of patients and extreme negative deviations in 4.97% (N=57) of patients (Supplementary Fig. 7a). From the perspective of brain regions, 27.73% (N=61) of the nodes showed an extreme FCS deviation in at least one patient, including extreme positive deviations in 3.64% (N=8) of brain regions and extreme negative deviations in 24.55% (N=54) of brain regions. Extreme positive deviations were found in the prefrontal cortex, angular gyrus, and subcortical areas, and extreme negative deviations were mainly found in the medial sensorimotor cortex and the temporal lobe (Supplementary Fig. 7b). The percentage of patients who extremely deviated from the normative range was remarkably low in either positive (≤0.26%) or negative (≤0.78%) deviations. We found that the brain regions with extreme deviations were slightly sparser when FDR was performed. However, we can also deduce that the specific brain regions affected by MDD vary markedly among individual patients, which is consistent with our main results.

Supplementary Tables

**Supplementary Table 1.** Demographic and clinical characteristics of patients with MDD with treatment outcomes

| N=43 | Baseline | Follow up  (6-month) | *t*/*P* |
| --- | --- | --- | --- |
| Age, mean (SD), years | 34.02 (9.27) |  |  |
| Gender (M/F) | 18/25 |  |  |
| Duration of Illness, mean (SD), months | 40.35 (57.22) |  |  |
| Age at Illness Onset, mean (SD), years | 30.77 (8.77) |  |  |
| HDRS-24, mean (SD) | 33.51 (7.59) | 2.16 (2.49) | **26.34/<0.001** |

Abbreviations: SD, standard deviation; M, male; F, female; HDRS, Hamilton Depression Rating Scale. *t*/*p* values were determined by using paired sample t tests between baseline and follow-up.

**Supplementary Table 2.** Scan parameters of the R-fMRI data from each center

| Center | Scanner | TR (ms) | TE (ms) | FA (°) | FOV (mm^2^) | Matrix | Resolution (mm^2^) | Slices | Thickness (mm) | Gap (mm) | VOL |
| --- | --- | --- | --- | --- | --- | --- | --- | --- | --- | --- | --- |
| CMU | GE HDxT 3T | 2000 | 40 | 90 | 240×240 | 64×64 | 3.75×3.75 | 35 | 3 | 0 | 200 |
| CSU | GE HDxT 3T | 2000 | 30 | 90 | 220×220 | 64×64 | 3.44×3.44 | 33 | 4 | 0.6 | 180 |
| GCMU1 | GE HDxT 3T | 2000 | 30 | 90 | 220×220 | 64×64 | 3.44×3.44 | 36 | 3 | 1 | 185 |
| GCMU2 | GE HDxT 3T | 2000 | 30 | 90 | 240×240 | 64×64 | 3.75×3.75 | 33 | 4 | 0 | 250 |
| KMU | PHILIPS Achieva 3T | 2200 | 35 | 90 | 230×230 | 128×128 | 1.80×1.80 | 50 | 3 | 0 | 240 |
| PKU | Siemens Trio 3T | 2000 | 30 | 90 | 210×210 | 64×64 | 3.28×3.28 | 30 | 4 | 0.8 | 210 |
| SCU | GE EXCITE 3T | 2000 | 30 | 90 | 220×220 | 64×64 | 3.44×3.44 | 30 | 5 | 0 | 200 |
| SWU | Siemens Trio 3T | 2000 | 30 | 90 | 220×220 | 64×64 | 3.44×3.44 | 32 | 3 | 1 | 242 |
| YMU | Siemens Trio 3T | 2500 | 27 | 77 | 220×220 | 64×64 | 3.44×3.44 | 43 | 3.4 | 0 | 200 |
| ZZU | GE MR750 3T | 2000 | 40 | 90 | 220×220 | 64×64 | 3.44×3.44 | 32 | 4 | 0.5 | 180 |

Abbreviations: TR, repetition time; TE, echo time; FA, flip angle; FOV, field of view; VOL, volume; CMU, China Medical University; CSU, Central South University; GCMU, Guangzhou University of Chinese Medicine; KMU, Kunming Medical University; PKU, Peking University; SCU, Sichuan University; SWU, Southwest University; YMU, National Yang-Ming University; ZZU, Zhengzhou University; GE, General Electric.

**Supplementary Table 3.** Clusters with significant group differences in the individual deviation maps between MDD patients and HCs

| No. | Region | Cluster Size | *P* value | Peak Coordinate | | | | |
| --- | --- | --- | --- | --- | --- | --- | --- | --- |
|  |  |  |  | X | | Y | | Z |
| **MDD > HCs** | | | | | | | | |
| 1 | Bilateral Precuneus/Angular/Inferior Parietal Gyrus/Superior Parietal Gyrus/Supramarginal Gyrus, BA 7/31/39/40 | 2394 | **<0.005** | -48 | -69 | | 30 | |
| 2 | Left Middle Frontal Gyrus/Superior Frontal Gyrus/Inferior Frontal Gyrus, BA 6/8/9 | 1297 | **<0.006** | -27 | 15 | | 42 | |
| 3 | Right Middle Frontal Gyrus/Superior Frontal Gyrus/Inferior Frontal Gyrus, BA 8/9/10 | 1044 | **<0.006** | 42 | 12 | | 24 | |
| 4 | Bilateral Thalamus | 237 | **<0.002** | 18 | -33 | | 6 | |
| 5 | Left Middle Temporal Gyrus/Inferior Temporal Gyrus, BA 20/21 | 208 | **<0.004** | -57 | -36 | | -27 | |
| 6 | Left Hippocampus | 191 | **<0.003** | -33 | -36 | | -12 | |
| **MDD < HCs** | | | | | | | | |
| 1 | Left ParaHippocampal Gyrus/Fusiform/Amygdala/Temporal pole, BA 28/34/36 | 344 | **<0.001** | -24 | -9 | | -39 | |
| 2 | Right Rolandic operculum/Insula/Superior Temporal Gyrus/Heschl Gyrus, BA 13/41 | 264 | **<0.002** | 48 | -27 | | 6 | |
| 3 | Right Inferior Occipital Gyrus/Lingual Gyrus, BA18 | 120 | **<0.005** | 24 | -99 | | -15 | |
| 4 | Right Median Cingulate and Paracingulate Gyri, BA 24 | 80 | **<0.007** | 3 | 3 | | 30 | |
| 5 | Left Paracentral lobule/Precentral, BA 6 | 51 | **<0.002** | -21 | -21 | | 60 | |

**Supplementary Table 4.** Network-level differences in mean deviation values between MDD subtypes

|  | VIS | SMN | DAN | VAN | LIM | FPN | DMN | SUB |
| --- | --- | --- | --- | --- | --- | --- | --- | --- |
| Subtype 1, mean (SD) | -0.02 (0.46) | -0.48 (0.46) | -0.26 (0.49) | -0.26 (0.48) | 0.12 (0.45) | 0.11 (0.58) | 0.32 (0.32) | 0.14 (0.56) |
| Subtype 2, mean (SD) | -0.02 (0.46) | 0.19 (0.36) | 0.19 (0.42) | 0.16 (0.44) | -0.12 (0.43) | 0.06 (0.50) | -0.15 (0.29) | -0.05 (0.58) |
| Subtype 1 vs. subtype 2 (*t/p*) | -0.14/0.892 | -27.03/<0.001 | -16.18/<0.001 | -15.09/<0.001 | 9.06/<0.001 | 1.32/0.187 | 25.56/<0.001 | 5.27/<0.001 |

Abbreviations: VIS, visual network; SMN, sensorimotor network; DAN, dorsal attention network; VAN, ventral attention network; LIB, limbic network; FPN, frontoparietal network; DMN, default mode network; SUB, subcortical regions.

**Supplementary Table 5.** Deviation differences among MDD subtypes and HCs

|  | The number of extremely deviated regions | The sum of positive extreme deviations | The sum of negative extreme deviations |
| --- | --- | --- | --- |
| Three Group (*F/p*) | 100.13/<0.001 | 55.99/<0.001 | 59.31/<0.001 |
| Subtype 1 vs. HCs (*t/p*) | 11.53/<0.001 | 8.90/<0.001 | -8.72/<0.001 |
| Subtype 2 vs. HCs (*t/p*) | -2.72/0.007 | -1.07/0.283 | 2.61/0.009 |
| Subtype 1 vs. subtype 2 (*t/p*) | 12.78/<0.001 | 8.49/<0.001 | -10.24/<0.001 |

**Supplementary Table 6.** Demographic and clinical differences between MDD subtypes

|  | Subtype 1 | Subtype 2 | *t* or *χ^2^*/*P* |
| --- | --- | --- | --- |
| Age | 35.35±16.05 (N=425) | 32.94±14.24 (N=723) | **2.64/0.008** |
| Gender(M/F) | 179/246 | 296/427 | 0.15/0.696 |
| HDRS total score | 21.13±6.92 (N=394) | 21.41±6.68 (N=671) | -0.65/0.515 |
| Age at Illness Onset | 33.53±13.82 (N=142) | 32.23±11.35 (N=221) | 0.98/0.330 |
| Duration | 2.24±3.74 (N=390) | 2.02±3.51 (N=669) | 0.94/0.349 |
| Episode (First/Recurrent) | 185/30 | 327/49 | 0.10/0.751 |
| Medicated (Yes/No) | 117/211 | 158/411 | **6.11/0.013** |
| 17 items of HDRS | (N=187) | (N=299) |  |
| Depressed Mood | 2.41±1.21 | 2.67±1.09 | **-2.42/0.016** |
| Feelings of Guilt | 1.05±0.93 | 1.05±0.99 | 0.04/0.971 |
| Suicide | 1.33±1.15 | 1.11±1.15 | **2.02/0.044** |
| Insomnia Early | 1.28±0.91 | 1.32±0.88 | -0.48/0.633 |
| Insomnia Middle | 1.19±0.83 | 1.31±0.80 | -1.69/0.09 |
| Insomnia Late | 1.04±0.87 | 1.11±0.87 | -0.79/0.429 |
| Work and Activities | 2.01±1.18 | 2.33±1.10 | **-3.11/0.002** |
| Retardation | 1.33±1.08 | 1.45±1.12 | -1.13/0.259 |
| Agitation | 0.77±0.92 | 0.91±1.00 | -1.54/0.123 |
| Anxiety Psychic | 1.70±1.20 | 1.90±1.07 | -1.94/0.053 |
| Anxiety, Somatic | 1.66±0.96 | 1.67±1.02 | -0.16/0.876 |
| Somatic Symptoms Gastrointestinal | 0.65±0.67 | 0.74±0.76 | -1.32/0.189 |
| Somatic Symptoms General | 0.95±0.76 | 0.86±0.81 | 1.21/0.227 |
| Genital Symptoms | 0.54±0.80 | 0.44±0.73 | 1.44/0.151 |
| Hypochondriasis | 0.83±0.97 | 0.71±0.90 | 1.38/0.167 |
| Loss Of Weight Within the Last Week | 0.36±0.66 | 0.29±0.62 | 1.23/0.220 |
| Insight | 0.43±0.61 | 0.51±0.67 | -1.30/0.194 |

Note: Data are presented as the mean±SD (number). M, male; F, female; HDRS, Hamilton Depression Rating Scale.

**Supplementary Table 7.** Subtype differences in the association between HDRS-17 total score and the onset age

| Source | df | Sum of Squares | Mean Square | *F* value | *P* value |
| --- | --- | --- | --- | --- | --- |
| Subtype | 1 | 156.68 | 156.68 | 2.25 | 0.135 |
| Onset age | 1 | 303.64 | 303.64 | 4.36 | **0.038** |
| Subtype*Onset Age | 1 | 307.05 | 307.05 | 4.41 | **0.037** |
| Error | 352 | 0.25×10^5^ | 69.64 | - | - |

**Supplementary Table 8.** Subtype differences in the association between HDRS-17 total score and the duration of illness

| Source | df | Sum of Squares | Mean Square | *F* value | *P* value |
| --- | --- | --- | --- | --- | --- |
| Subtype | 1 | 26.35 | 26.35 | 0.57 | 0.449 |
| Duration | 1 | 3.84 | 3.84 | 0.08 | 0.772 |
| Subtype*Duration | 1 | 174.04 | 174.04 | 3.79 | 0.052 |
| Error | 1006 | 0.46×10^5^ | 45.87 | - | - |

**Supplementary Table 9.** Demographic and clinical differences between MDD subtypes in leave-one-site-out validation.

| Leave-out Site | CMU | CSU | GCMU1 | GCMU2 | KMU | PKU | SCU | SWU | YMU | ZZU |
| --- | --- | --- | --- | --- | --- | --- | --- | --- | --- | --- |
| Age | **3.61/0.000** | **2.19/0.029** | **2.84/0.005** | **2.95/0.003** | **2.63/0.009** | **3.00/0.003** | **2.52/0.012** | **3.41/0.001** | 1.52/0.129 | 1.92/0.055 |
| Gender(M/F) | 0.02/0.894 | 0.03/0.860 | 0.00/0.998 | 0.65/0.419 | 0.12/0.731 | 0.02/0.903 | 0.13/0.714 | 0.06/0.814 | 0.86/0.354 | 0.24/0.622 |
| HDRS total score | -1.51/0.131 | -0.30/0.768 | -0.88/0.379 | -0.37/0.712 | -0.68/0.499 | -0.49/0.622 | -0.34/0.731 | -1.26/0.210 | 0.47/0.641 | -0.41/0.684 |
| Onset Age | **2.50/0.013** | 0.26/0.797 | -0.99/0.321 | -1.06/0.290 | -1.24/0.214 | -1.51/0.132 | -0.41/0.684 | -1.19/0.237 | 1.05/0.293 | 1.15/0.252 |
| Duration | -0.60/0.547 | 0.53/0.600 | 0.90/0.366 | 0.59/0.554 | 0.95/0.340 | 1.03/0.303 | 1.45/0.148 | 1.79/0.074 | **1.99/0.047** | 0.78/0.434 |
| Episode (First/Recurrent) | 0.17/0.683 | 0.05/0.833 | 0.24/0.628 | 0.08/0.779 | 0.15/0.700 | 0.52/0.470 | 0.69/0.407 | **4.00/0.046** | 0.02/0.885 | 0.24/0.626 |
| Medicated (Yes/No) | **6.23/0.013** | **4.24/0.039** | **7.16/0.007** | 3.05/0.081 | **6.08/0.014** | **6.59/0.010** | 3.81/0.051 | **8.88/0.003** | 1.62/0.203 | 2.54/0.111 |
| 17 items of HDRS |  | | | | | | | | | |
| Depressed Mood | **-2.23/0.026** | **-2.31/0.022** | **-2.75/0.006** | **-2.16/0.032** | **-2.41/0.016** | **-2.03/0.043** | **-2.53/0.012** | **-2.59/0.010** | -1.49/0.136 | -1.52/0.130 |
| Feelings of Guilt | -0.51/0.607 | -0.22/0.824 | 0.04/0.971 | 0.25/0.803 | 0.14/0.886 | 0.20/0.841 | -0.10/0.922 | -0.41/0.680 | -0.26/0.795 | -0.16/0.873 |
| Suicide | 1.80/0.073 | 1.61/0.108 | 1.85/0.065 | **2.01/0.045** | **1.97/0.049** | **2.98/0.003** | 1.71/0.087 | 0.45/0.652 | 0.99/0.325 | 0.29/0.770 |
| Insomnia Early | 0.13/0.895 | 0.02/0.983 | -0.48/0.633 | 0.07/0.946 | -0.21/0.838 | -0.61/0.542 | -0.21/0.837 | -1.31/0.191 | 0.28/0.779 | 0.68/0.497 |
| Insomnia Middle | -1.63/0.104 | -1.52/0.129 | -1.69/0.092 | -1.69/0.092 | -1.86/0.063 | **-2.14/0.033** | -1.80/0.072 | **-2.29/0.023** | -0.88/0.382 | -0.80/0.426 |
| Insomnia Late | -0.20/0.838 | -0.27/0.788 | -0.79/0.429 | -0.44/0.663 | -0.67/0.504 | -0.54/0.589 | -0.64/0.520 | -1.37/0.171 | 0.18/0.860 | -0.54/0.593 |
| Work and Activities | **-3.45/0.001** | **-2.65/0.008** | **-3.28/0.001** | **-3.22/0.001** | **-3.25/0.001** | **-3.46/0.001** | **-2.73/0.007** | **-2.64/0.009** | -1.72/0.086 | **-2.72/0.007** |
| Retardation | -1.61/0.108 | -1.34/0.182 | -1.30/0.195 | -1.25/0.211 | -1.23/0.218 | -0.98/0.328 | -1.22/0.222 | -1.22/0.223 | -0.20/0.843 | -0.79/0.430 |
| Agitation | -1.93/0.054 | -1.14/0.254 | -1.74/0.083 | -1.41/0.159 | -1.57/0.116 | -1.59/0.112 | -1.42/0.156 | -1.89/0.060 | -1.59/0.113 | -0.50/0.621 |
| Anxiety Psychic | **-2.46/0.014** | -1.55/0.122 | **-2.11/0.036** | **-2.09/0.037** | **-1.97/0.049** | -1.97/0.050 | -1.48/0.141 | -1.67/0.095 | -0.05/0.958 | -1.32/0.189 |
| Anxiety, Somatic | -0.72/0.469 | 0.00/1.000 | -0.25/0.803 | -0.38/0.706 | -0.31/0.754 | -0.08/0.933 | -0.37/0.710 | 0.22/0.829 | 0.12/0.906 | 0.12/0.904 |
| Somatic Symptoms Gastrointestinal | -1.17/0.244 | -0.90/0.371 | -1.45/0.149 | -1.22/0.224 | -1.14/0.256 | -1.12/0.263 | -1.26/0.209 | -1.51/0.133 | -1.33/0.183 | 0.18/0.855 |
| Somatic Symptoms General | 0.43/0.668 | 0.43/0.669 | 1.09/0.276 | 1.16/0.245 | 1.19/0.236 | 1.28/0.200 | 0.71/0.478 | 0.63/0.528 | 1.45/0.148 | -0.68/0.496 |
| Genital Symptoms | 0.57/0.572 | 0.97/0.331 | 1.44/0.151 | 1.19/0.236 | 1.56/0.120 | **2.11/0.035** | 0.84/0.401 | -0.36/0.721 | 1.25/0.210 | 0.25/0.802 |
| Hypochondriasis | 1.34/0.180 | 1.71/0.088 | 1.38/0.167 | 1.49/0.136 | 1.54/0.125 | 1.80/0.073 | 1.77/0.078 | 1.32/0.186 | **2.32/0.021** | 0.39/0.700 |
| Loss Of Weight Within the Last Week | 0.34/0.737 | 1.58/0.114 | 1.23/0.220 | 0.83/0.407 | 1.03/0.304 | 0.93/0.355 | 1.43/0.153 | -0.47/0.639 | 1.25/0.211 | 0.99/0.322 |
| Insight | **-2.06/0.040** | -1.11/0.269 | -1.45/0.149 | -1.75/0.080 | -1.46/0.146 | -1.31/0.192 | -1.41/0.158 | -1.64/0.102 | -1.49/0.138 | -0.09/0.936 |
| Association between HDRS score and onset age |  | | | | | | | | | |
| subtype*onset age | 1.02/0.314 | **3.98/0.047** | **4.51/0.034** | 3.17/0.076 | **4.21/0.041** | 1.55/0.214 | **4.41/0.037** | **8.14/0.005** | 2.85/0.092 | 3.45/0.064 |
| post-hoc (subtype1) | - | **-0.30/0.001** | **-0.24/0.004-** | **-** | **-0.24/0.005** | **-** | **-0.28/0.001** | **-0.26/<0.001** | **-** | **-** |
| post-hoc (subtype2) | - | -0.04/0.613 | 0.00/0.985 | - | -0.01/0.932 | - | -0.03/0.716 | 0.05/0.493 | - | - |
| Prediction of treatment response |  | | | | | | | | | |
| Correlation between predicted and observed HDRS change (subtype 1) | -0.01/0.476 | **-** | **0.47/0.014** | **0.50/0.007** | **0.50/0.010** | 0.34/0.053 | 0.15/0.263 | 0.17/0.234 | 0.15/0.247 | **0.50/0.016** |
| Correlation between predicted and observed HDRS change (subtype 2) | -0.28/0.958 | - | -0.14/0.782 | -0.30/0.934 | -0.30/0.940 | -0.18/0.821 | -0.15/0.796 | -0.15/0.718 | -0.15/0.786 | -0.30/0.953 |

Note: Data are presented as *t*, *χ^2^, F,* or *r*/*P*. M, male; F, female; HDRS, Hamilton Depression Rating Scale; CMU, China Medical University; CSU, Central South University; GCMU, Guangzhou University of Chinese Medicine; KMU, Kunming Medical University; PKU, Peking University; SCU, Sichuan University; SWU, Southwest University; YMU, National Yang-Ming University; ZZU, Zhengzhou University.

Supplementary Figures

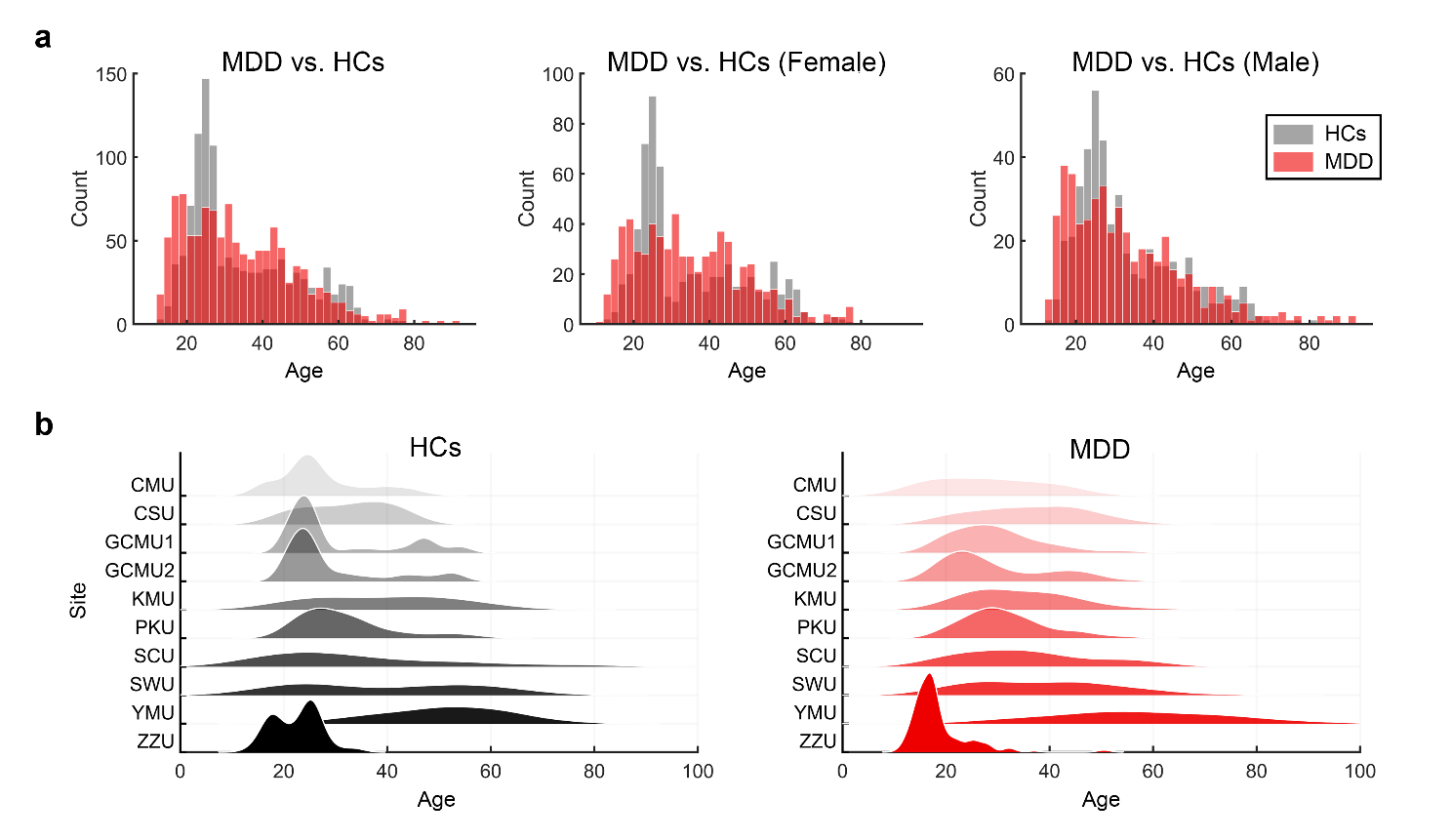

**Supplementary Fig. 1** Age distribution of participants. **a** Age distribution (x-axis) of all patients with MDD and HCs and the gender-specific distributions. **b** Age distribution (x-axis) of each site (y-axis) in the HCs and patients with MDD. MDD, major depressive disorder; HCs: healthy controls; CMU, China Medical University; CSU, Central South University; GCMU, Guangzhou University of Chinese Medicine; KMU, Kunming Medical University; PKU, Peking University; SCU, Sichuan University; SWU, Southwest University; YMU, National Yang-Ming University; ZZU, Zhengzhou University.

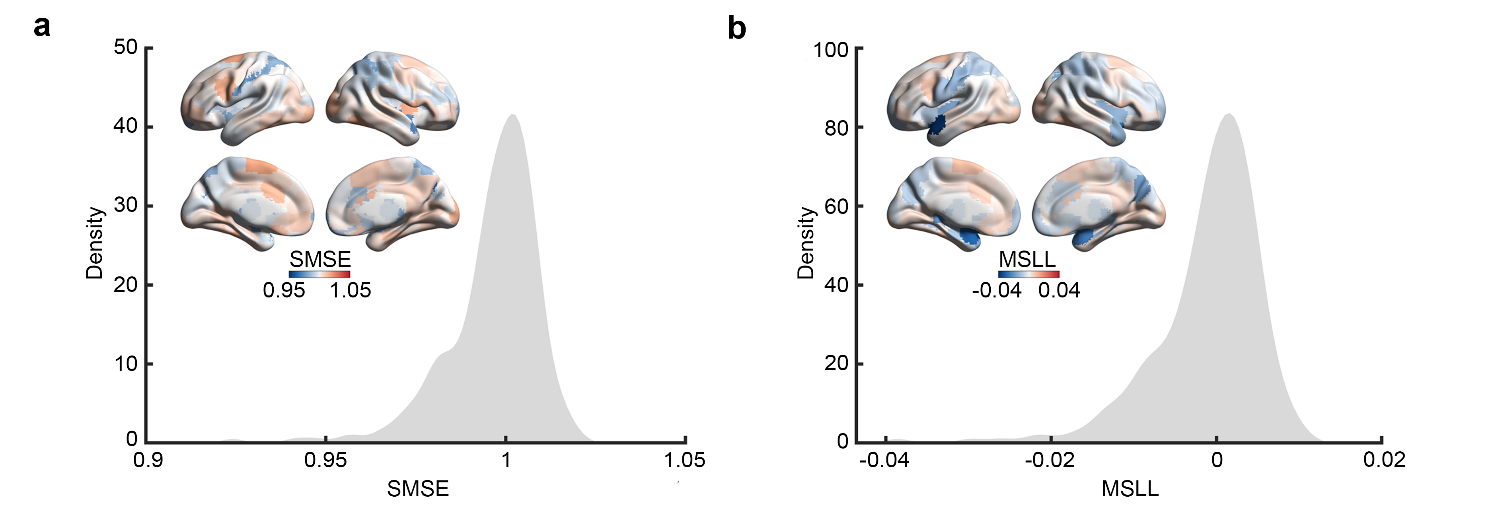

**Supplementary Fig. 2** Model fit evaluation. The distribution and brain map of SMSE (**a**) and MSLL (**b**) between observed and predicted FCS values in HCs under 10-fold cross-validation. SMSE, standardized mean squared error; MSLL, mean squared log-loss; FCS, functional connectivity strength.

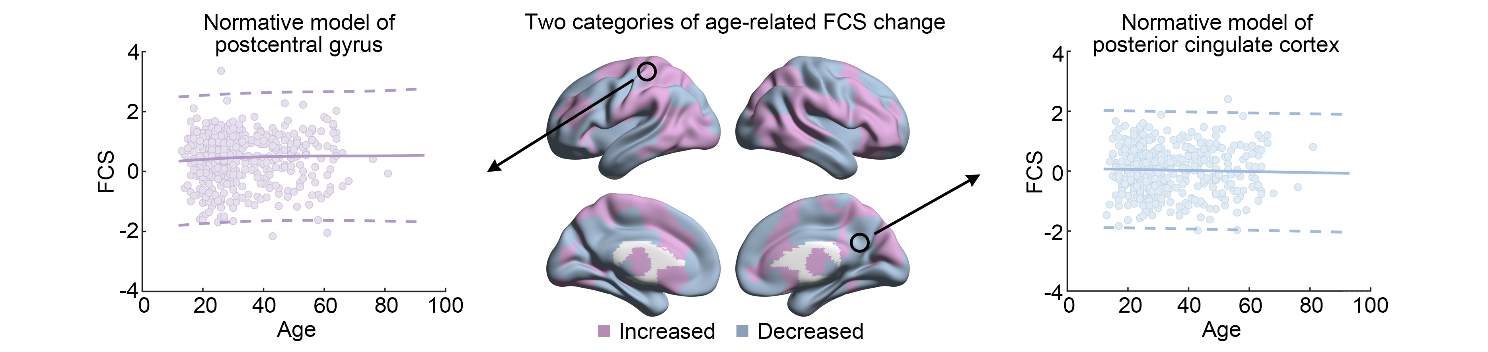

**Supplementary Fig. 3** Normative models established in HCs. The brain map in the middle indicates the two categories of age-related FCS change trajectories (purple: increased; blue: decreased) in HCs (male). The FCS change trajectories (solid line) and the normative range (dashed line) of postcentral gyrus and posterior cingulate cortex are shown on the left and right. Each dot represents the data from one HC. HCs, healthy controls; FCS, functional connectivity strengths.

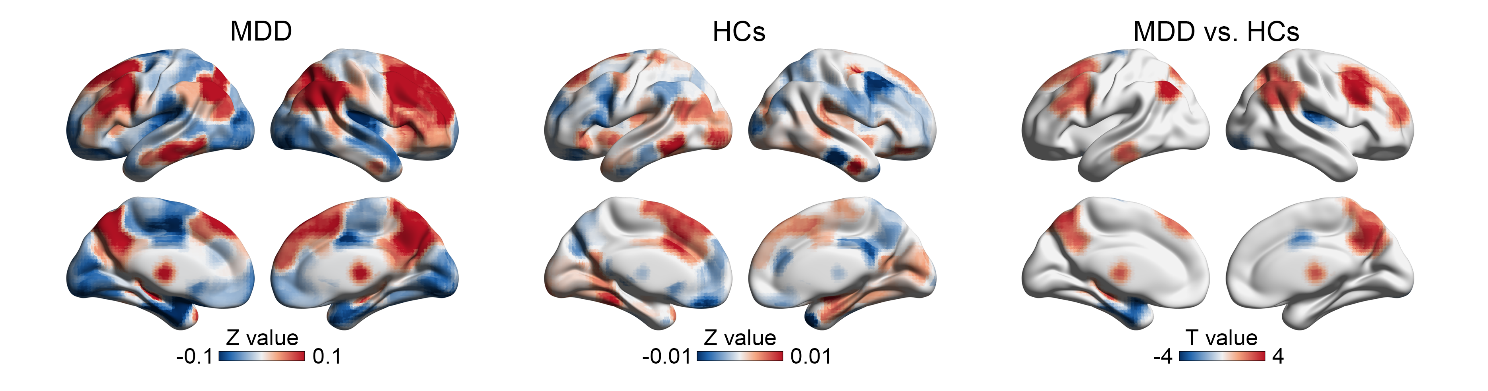

**Supplementary Fig. 4** Mean deviation map of MDD patients and HCs, and their group differences. HCs, healthy controls; MDD, major depressive disorder.

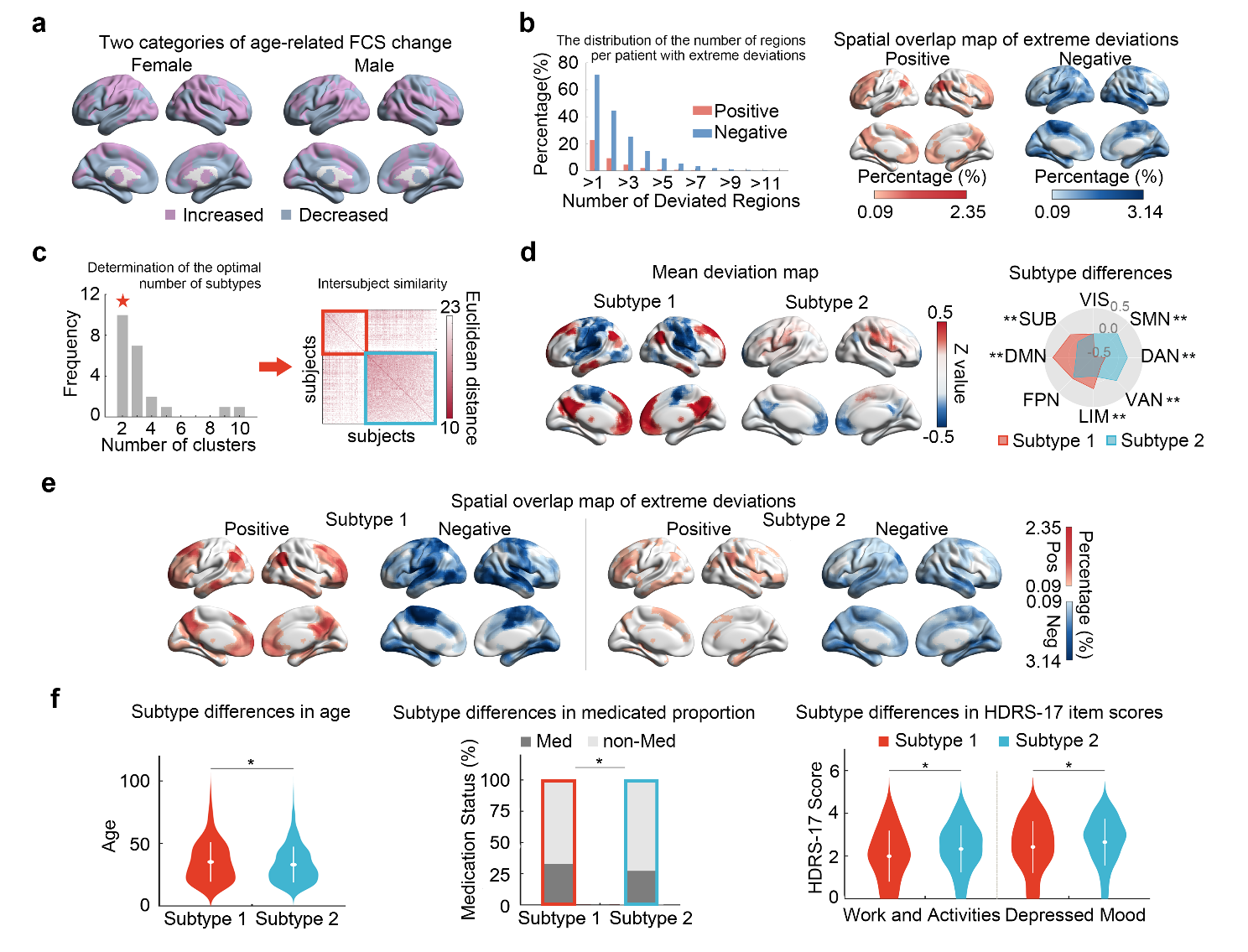

**Supplementary Fig. 5** The validation results under different thresholds in the FCS calculation (r=0.15). **a** The brain regions were clustered into two categories according to their age-related FCS change trajectories in both female (left) and male (right) groups. **b** Bar plots show the distribution of the number of regions per patient with extremely positive (red) and negative (blue) deviations. The spatial overlap maps indicate the percentage of patients who extremely deviated from the normative range for each brain region (left, extreme positive deviations; right, extreme negative deviations). **c** Determination of the optimal number of MDD subtypes and the intersubject differences in the FCS deviation patterns among patients. **d** The mean deviation map of each subtype and their system-level differences. **e** The spatial overlap map of extreme positive and negative deviations of each subtype. **f** Subtype differences in demographic and clinical variables. VIS, visual network; SMN, sensorimotor network; DAN, dorsal attention network; VAN, ventral attention network; LIB, limbic network; FPN, frontoparietal network; DMN, default mode network; SUB, subcortical regions; HDRS, Hamilton Depression Rating Scale; FCS, functional connectivity strength; MDD, major depressive disorder; ^*^*p*<0.05; ^**^*p*<0.05, FDR corrected.

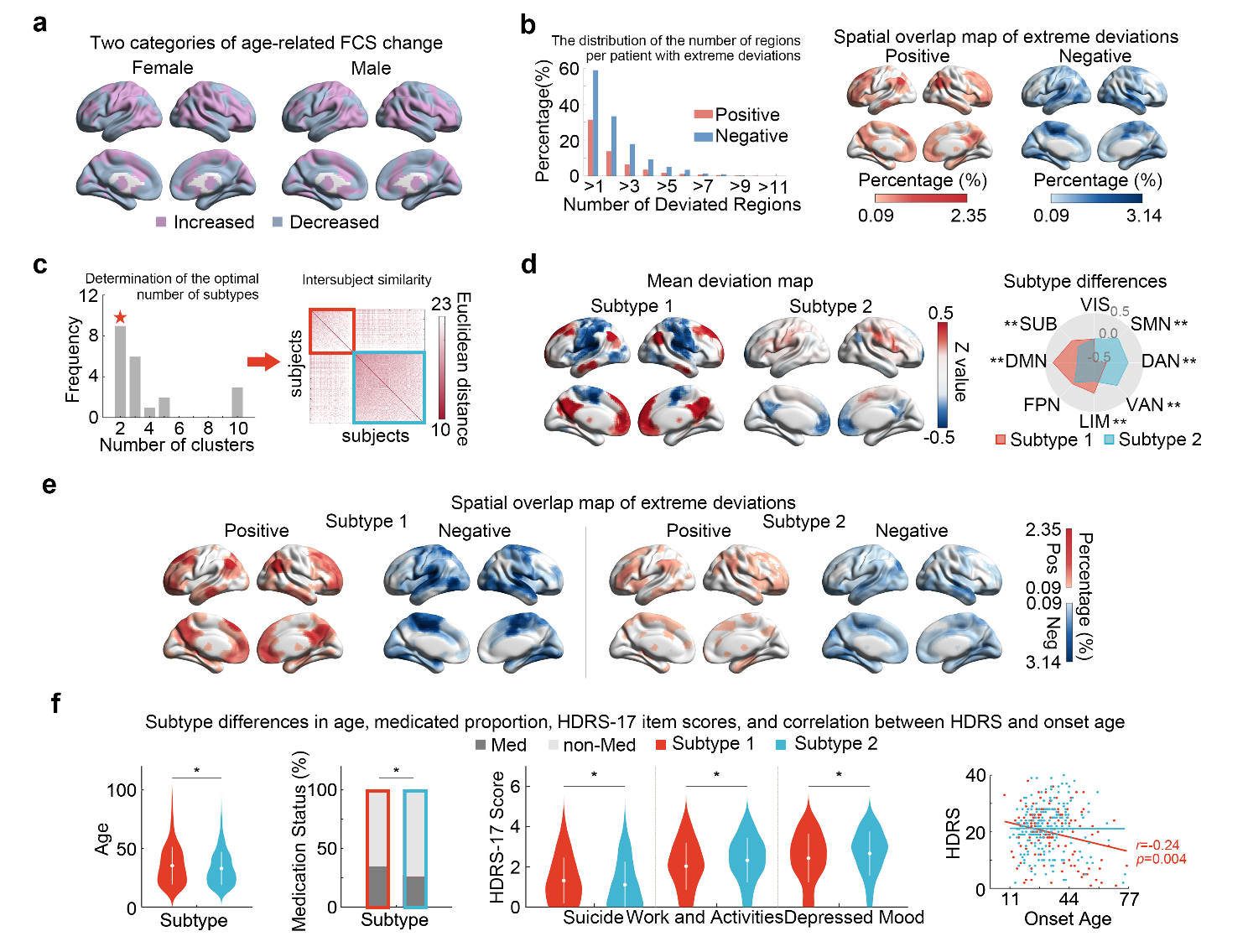

**Supplementary Fig. 6** The validation results under different thresholds in the FCS calculation (r=0.25). **a** The brain regions were clustered into two categories according to their age-related FCS change trajectories in both female (left) and male (right) groups. **b** Bar plots show the distribution of the number of regions per patient with extremely positive (red) and negative (blue) deviations. The spatial overlap maps indicate the percentage of patients who deviated extremely from the normative range for each brain region (left, extreme positive deviations; right, extreme negative deviations). **c** Determination of the optimal number of MDD subtypes and the intersubject differences in the FCS deviation patterns among patients. **d** The mean deviation map of each subtype and their system-level differences. **e** The spatial overlap map of extreme positive and negative deviations of each subtype. **f** Subtype differences in demographic and clinical variables. VIS, visual network; SMN, sensorimotor network; DAN, dorsal attention network; VAN, ventral attention network; LIB, limbic network; FPN, frontoparietal network; DMN, default mode network; SUB, subcortical regions; HDRS, Hamilton Depression Rating Scale; FCS, functional connectivity strength; MDD, major depressive disorder; ^*^*p*<0.05; ^**^*p*<0.05, FDR corrected.

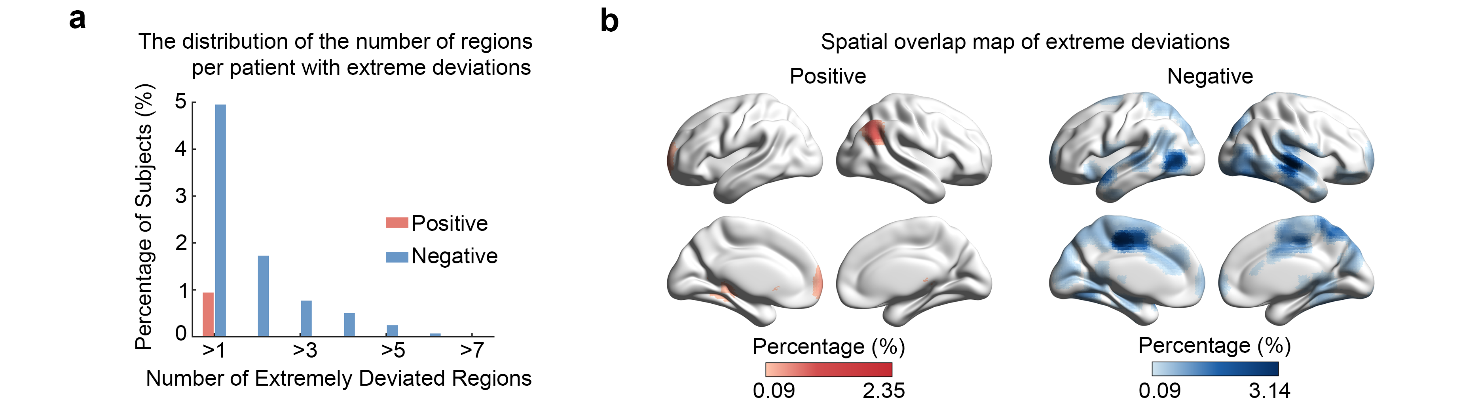

**Supplementary Fig. 7** Characterization of extreme deviations from the normative model using FDR *p*<0.05. **a** Bar plots show the distribution of the number of regions per patient with extremely positive (red) and negative (blue) deviations. **b** The spatial overlap maps indicate the percentage of patients who deviated extremely from the normative range for each brain region (left, extreme positive deviations; right, extreme negative deviations).

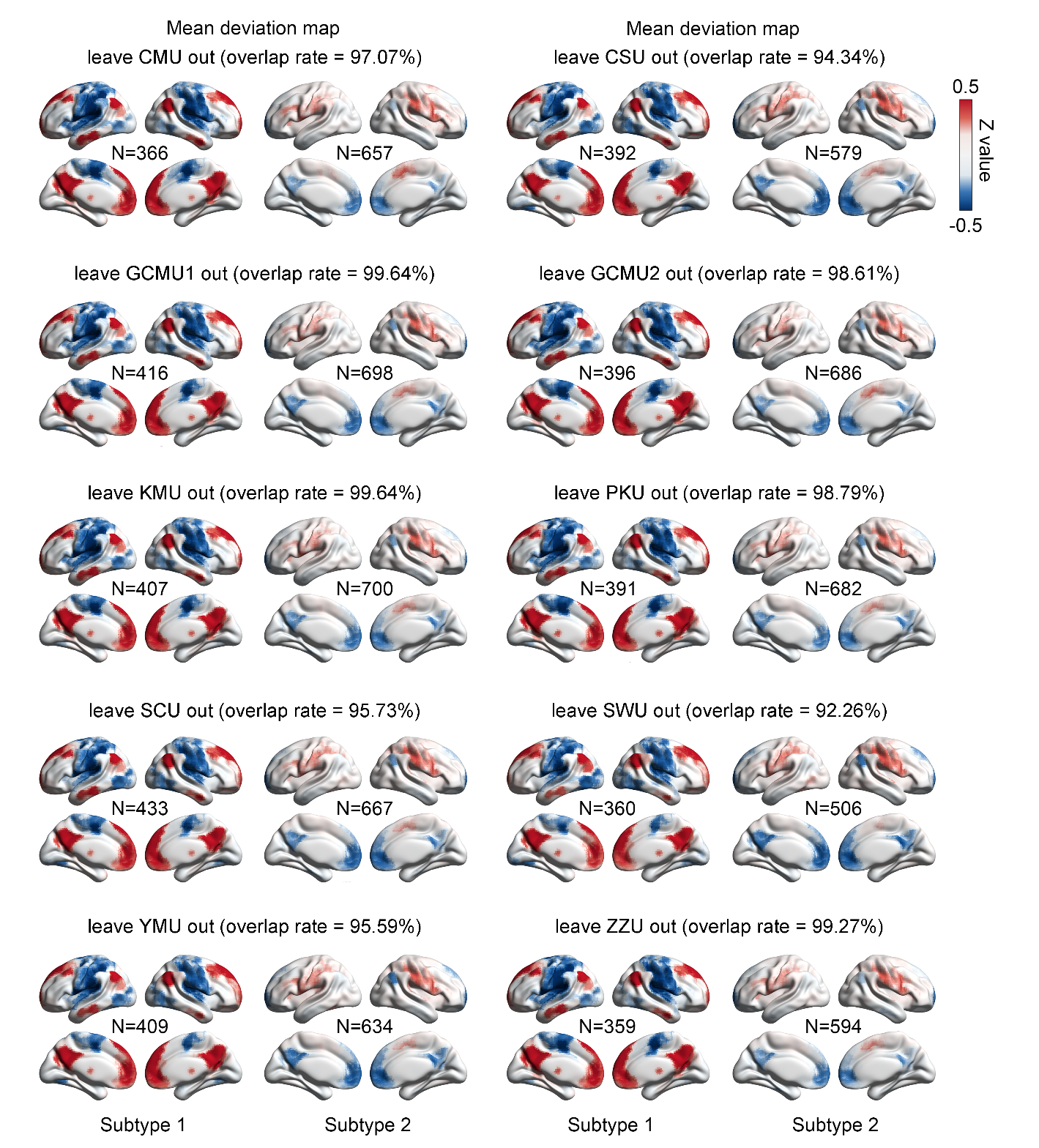

**Supplementary Fig. 8** The mean deviation map of each subtype when performing leave-one-site-out validation. The overlap rates of the resulting clustered indexes with the clustered indexes in the main results were all >92%. CMU, China Medical University; CSU, Central South University; GCMU, Guangzhou University of Chinese Medicine; KMU, Kunming Medical University; PKU, Peking University; SCU, Sichuan University; SWU, Southwest University; YMU, National Yang-Ming University; ZZU, Zhengzhou University.
